# The *Drosophila* histone demethylase KDM5 is required during early neurodevelopment for proper mushroom body formation and cognitive function

**DOI:** 10.1101/2020.10.09.324939

**Authors:** Hayden A. M. Hatch, Helen M. Belalcazar, Owen J. Marshall, Julie Secombe

## Abstract

Mutations in the lysine demethylase 5 (KDM5) family of transcriptional regulators are associated with intellectual disability, yet little is known regarding the spatiotemporal requirements or neurodevelopmental contributions of KDM5 proteins. Utilizing the mushroom body (MB), a major learning and memory center within the *Drosophila* brain, we demonstrate that KDM5 is specifically required within ganglion mother cells and immature neurons for proper neurodevelopment and cognitive function. Within this cellular subpopulation, we identify a core network of KDM5-regulated genes that are critical modulators of neurodevelopment. Significantly, we find that a majority of these genes are direct targets of Prospero (Pros), a transcription factor with well-established roles in neurodevelopment in other neuronal contexts. We demonstrate that Pros is essential for MB development and functions downstream of KDM5 to regulate MB morphology. We therefore provide evidence for a KDM5-Pros axis that orchestrates a transcriptional program critical for proper axonal development and cognitive function.

## INTRODUCTION

Intellectual disability (ID) is reported to affect 1.5% to 3% of the global population and represents a class of neurodevelopmental disorders characterized by cognitive impairments that result in lifelong educational, social, and financial consequences for patients and their caregivers (van Bokhoven, 2011; Leonard and Wen, 2002). ID disorders are diagnosed during early childhood and are defined by an IQ score of less than 70 with deficits in adaptive behaviors (Ropers, 2010). However, despite these profound burdens, little is known regarding the pathogenesis of ID disorders, particularly how disruptions in genetic and neuronal regulatory programs contribute to cognitive and behavioral dysfunction.

Advances in comparative genomic hybridization and whole exome sequencing have revealed strong associations between ID and mutations in genes encoding chromatin modifying enzymes (Elbert and Bérubé, 2014; Harmeyer et al., 2017; Kong et al., 2017; Parkel et al., 2013). These proteins post-translationally modify chromatin by inserting or removing chemical moieties from histone tails to regulate DNA accessibility (van Bokhoven, 2011; Vallianatos and Iwase, 2015). They may also act through the recruitment of other proteins to modulate transcriptional initiation or elongation (Liefke et al., 2010; Liu et al., 2014; Secombe et al., 2007; van Oevelen et al., 2008). One class of chromatin modifiers is the Lysine Demethylase 5 (KDM5) family of transcriptional regulators, with mammals encoding four *KDM5* paralogs: *KDM5A*, *KDM5B*, *KDM5C*, and *KDM5D.* Mutations in *KDM5A*, *KDM5B*, and *KDM5C* are associated with ID, with genetic variants in *KDM5C* associated with a disorder known as Mental Retardation, X-linked, Syndromic, Claes-Jensen type (MRXSCJ, OMIM# 300534). Notably, mutations in *KDM5C* are predicted to account for 0.7-2.8% of all X-linked ID, ranging from mild to severe, and are associated with *KDM5C* loss of function (Gonçalves et al., 2014; Ropers & Hamel, 2005).

The generation of *KDM5* knockout animal models has greatly assisted in our ability to investigate the neuromorphological and behavioral consequences of *KDM5* loss of function. Previous *in vitro* studies examining rat cerebellar granular neurons and pyramidal neurons of prepared mouse basolateral amygdala slices demonstrate that *Kdm5c* knockout results in dendritic spine abnormalities (Iwase, Brookes, and Agarwal et al., 2016). Additionally, *Kdm5c* knockout mice display behavioral deficits that are analogous to those exhibited by MRXSCJ patients, such as increased aggression, learning and memory impairments, and decreased seizure thresholds (Iwase, Brookes, and Agarwal et al., 2016; Scandaglia et al., 2017). Similarly, loss of function mutations in *rbr-2*, the sole *C. elegans Kdm5c* ortholog, result in axonal growth and guidance defects (Mariani et al., 2016). Together, these studies suggest that the neuromorphological and functional impairments resulting from loss of orthologous KDM5 proteins are likely to be attributed to altered gene expression within neurons.

KDM5 proteins demethylate trimethyl groups on lysine 4 of histone H3 (H3K4me3) *via* the enzymatic activity of their Jumonji C (JmjC) domains (Liefke et al., 2010; Liu et al., 2014; Secombe et al., 2007; van Oevelen et al., 2008). High levels of H3K4me3 near transcriptional start sites (TSS) are associated with actively transcribed genes, suggesting that KDM5 proteins can dynamically regulate transcription (Greer and Shi, 2012). Prevailing models linking alterations in KDM5 family protein function to ID suggest that loss of JmjC-mediated demethylase activity is a key driver of neuronal dysfunction (Mariani et al., 2016; Scandaglia et al., 2017; Vallianatos & Iwase, 2015; Vallianatos et al., 2020; Zamurrad et al., 2018). For example, Vallianatos and colleagues have demonstrated that the neuronal and behavioral phenotypes observed in *Kdm5c* knockout mice can be rescued by reducing levels of the H3K4 methyltransferase KMT2A (Vallianatos et al., 2020). Additionally, our lab has previously demonstrated that *Drosophila* KDM5 acts in a demethylase-dependent manner to regulate long- and short-term olfactory memory (Zamurrad et al., 2018).

KDM5 family proteins can also regulate transcription independent of their demethylase activity by associating with other chromatin modifying proteins (Gajan et al., 2016; M.G. Lee et al., 2007; N. Lee et al., 2009; Nishibuchi et al., 2014). For example, the HDAC complex member SIN3A has been shown *in vitro* to interact with *Drosophila* KDM5 and regulate overlapping subsets of genes. In mice, SIN3A is required for neuronal development (Gajan et al., 2016; Witteveen et al., 2016) with mutations in SIN3A associated with Witteveen-Kolk syndrome (OMIM# 613406), a neurodevelopmental disorder characterized by developmental delay and ID (Witteveen et al., 2016). Additionally, a subset of ID-associated *KDM5C* missense mutations have been shown *in vitro* not to affect H3K4me3 demethylase activity, yet alter transcriptional outputs (Brookes et al., 2015; Vallianatos et al., 2018). Collectively, these data provide strong evidence that disruption of KDM5 protein function may impact multiple transcriptional pathways critical to neuronal development and function.

Here, we utilize *Drosophila*, which encodes a single, highly conserved *KDM5* ortholog, known as *kdm5* or *little imaginal discs* (*lid*), to investigate the genetic, neuromorphological, and behavioral consequences of KDM5 loss during neurodevelopment. Our analyses focus on a group of neurons known as Kenyon cells, which form a bilateral, neuropil-rich structure known as the mushroom body (MB). The MB is essential for orchestrating a diverse repertoire of cognitive processes and is thus routinely used to study neuroanatomical changes associated with mutations in ID-related genes (Androschuk et al., 2015; Aso et al., 2014; Dubnau et al., 2001; Heisenberg et al., 1985). In fact, loss of function mutations in orthologous genes associated with ID, such as the Fragile X Syndrome gene *fmr1*, the Down Syndrome gene *dscam* and the *ZC3H14* autosomal recessive ID gene *dnab2*, result in severe morphological defects of the MB (Hattori et al., 2007; Kelly et al., 2016; Michel, 2004; Zhan et al., 2004).

The development of the MB is dependent on four MB neuroblasts (MBNBs) per hemisphere dividing asymmetrically throughout development. Each MBNB gives rise to another MBNB and a ganglion mother cell (GMC), which in turn divides symmetrically to form two Kenyon cells. Three subclasses of Kenyon cells give rise to the MB and are born in a highly regulated and sequential manner with tight temporal control. The first-born Kenyon cells are referred to as γ Kenyon cells and develop between the embryonic and mid-third instar larval stage, giving rise to the γ lobes. The α’/β’ lobes are the next to develop, followed by the α/β lobes during the late larval and pupal stages. Notably, specification of these Kenyon cell subsets are transcriptionally regulated though the timed expression of a number of transcription factors (Bates et al., 2010; Marchetti and Tavosanis, 2017; Syed et al., 2017).

We show here that *kdm5* knockout and shRNA-mediated depletion of *kdm5* within GMCs and immature neurons both result in profound defects to MB axonal guidance and growth. Cell-specific depletion of *kdm5* additionally results in significant behavioral deficits as revealed by altered avoidance response to *Drosophila* stress odorant (dSO). Furthermore, using an *in vivo* transcriptomics-based approach, we identify a subset of genes implicated in GMC fate determination and axon guidance that are downregulated by *kdm5* loss within GMCs and immature Kenyon cells. One such gene, *prospero* (*pros*), encodes a homeodomain-containing transcription factor with a critical role in promoting proper axon pathfinding and growth. While Pros plays well-established roles in other neuronal cell types, its importance in regulating MB development remains unexplored. We find here that MB-specific knockdown of *pros* leads to aberrant axonal growth and guidance. We additionally demonstrate that half of KDM5 regulated genes within GMCs and immature neurons are bound by Pros and that *kdm5* and *pros* lie within the same genetic pathway. Our studies thus provide the first *in vivo* analysis of KDM5 within a specific cell population, revealing a key genetic interaction between *kdm5* and *pros* critical for neurodevelopment.

## RESULTS

### KDM5 is expressed throughout the CNS and is essential for proper MB morphology

To assess the neurodevelopmental consequences resulting from KDM5 loss, we examined MB morphology of animals that were homozygous for a *kdm5* null allele, *kdm5*^*140*^ (Drelon et al., 2018, 2019). As homozygous *kdm5*^*140*^ animals fail to eclose from their pupal cases, we performed our immunohistochemical analysis on pharate adults, which externally appear indistinguishable from wild-type animals (Drelon et al., 2018). Following a well-established classification scheme used by others (Gombos et al., 2015; Kelly et al., 2016; Michel, 2004), MB defects were categorized as impacting MB growth and/or guidance, with the former defined by a stunted, overextended, or absent lobe and the latter by a misprojected lobe. Staining using an antibody specific to the NCAM-like cell adhesion molecule Fasciclin 2 (Fas2) revealed highly penetrant MB abnormalities, with ~70% of animals exhibiting growth and guidance defects of the α/β lobes (Fig. 1A,B). Because animals specifically lacking KDM5 demethylase activity have phenotypically normal α/β lobes (Zamurrad et al., 2018), these data demonstrate that KDM5 is required for maintaining MB morphology independent of its canonical enzymatic function.

**Figure 1.**
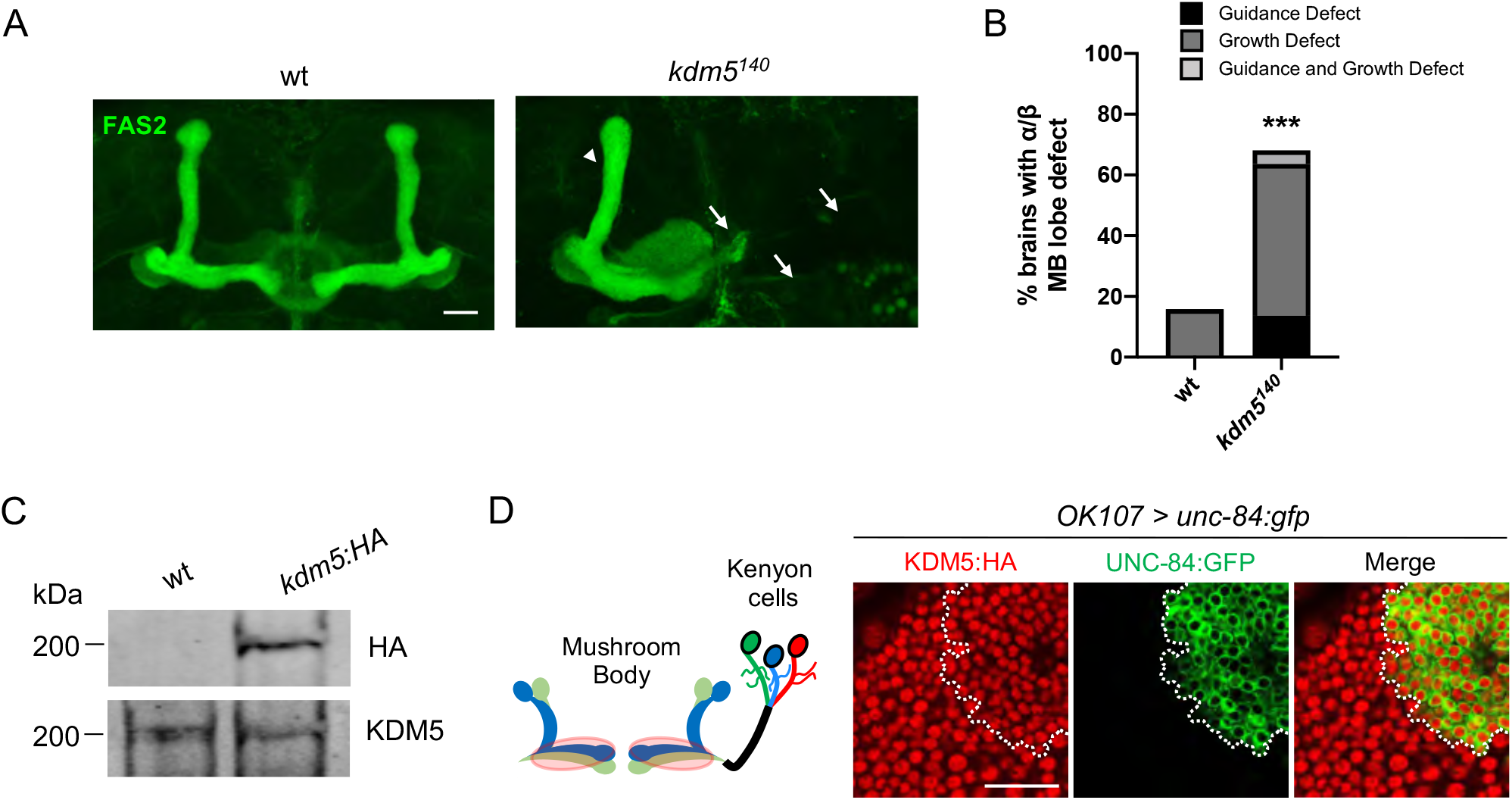
*kdm5*^*140*^ Pharate Adults have Neuromorphological Defects of the MB. (A) Representative α/β lobe Z projections of pharate wildtype (*NP4707*^*rev2*^ revertant) and *kdm5*^*140*^ strains. Arrows indicate growth defects and arrowheads indicate guidance defects. The α/β lobes are revealed with anti-Fas2. (B) Quantification of α/β lobe defects in wildtype and *kdm5*^*140*^ strains. *n* = 19-22 (mean *n* = 21). *** *P* < 0.001 (chi-square test with Yates’ correction). Scale bars represent 20 μm. (C) Western blot of *w*^*1118*^ and *kdm5:HA* adult heads confirming wild type expression levels of KDM5:HA within our endogenously-tagged *kdm5:HA* strain. Anti-HA (top) and anti-KDM5 (bottom) loading control. (D) Schematic of an adult MB with its associated Kenyon cells (left). *OK107-Gal4* is used to drive expression of UNC-84:GFP, an inner nuclear membrane GFP reporter, within MBNBs, MB-GMCs and Kenyon cells (right). Brains are counterstained with anti-HA to demonstrate the presence of endogenously tagged KDM5:HA within Kenyon cell nuclei.

To assist in our functional understanding of KDM5, we sought to investigate its expression throughout CNS development. To facilitate these analyses, we used CRISPR/Cas9 to generate a strain containing a *3xHA*-tag fused to the endogenous locus of *kdm5* and demonstrated by western blot that the KDM5:HA protein is expressed at the same level as endogenous KDM5 from wild-type animals (Fig. 1C). We also confirmed that KDM5:HA was expressed in adult MB Kenyon cells by co-expression of a nuclear membrane-localized GFP reporter, UNC-84:GFP, with the MB driver *OK107-Gal4* (Fig. 1D). Immunostaining of the central nervous system (CNS) of wandering third instar larvae revealed that KDM5 was localized to cortical nuclei while absent from neuropil-rich regions marked by the ubiquitous presynaptic active zone marker Bruchpilot (Brp) (Fig. 2A). KDM5:HA localized to cortical nuclei across a variety of cell types, including neurons (Fig. 2B), neuroblasts (NBs) and presumptive GMCs (Fig. 2C). We also examined KDM5:HA expression in the adult brain, where KDM5:HA appeared to be similarly localized to cortical nuclei while absent from neuropil-rich regions, such as the antennal lobes and both the dorsolateral and ventrolateral protocerebra (Fig. 2D,E). Given the broad expression pattern of KDM5:HA within a variety of cell types, KDM5 may regulate a range of neural processes, from NPC division and axonal growth to neuronal maturation and function.

**Figure 2.**
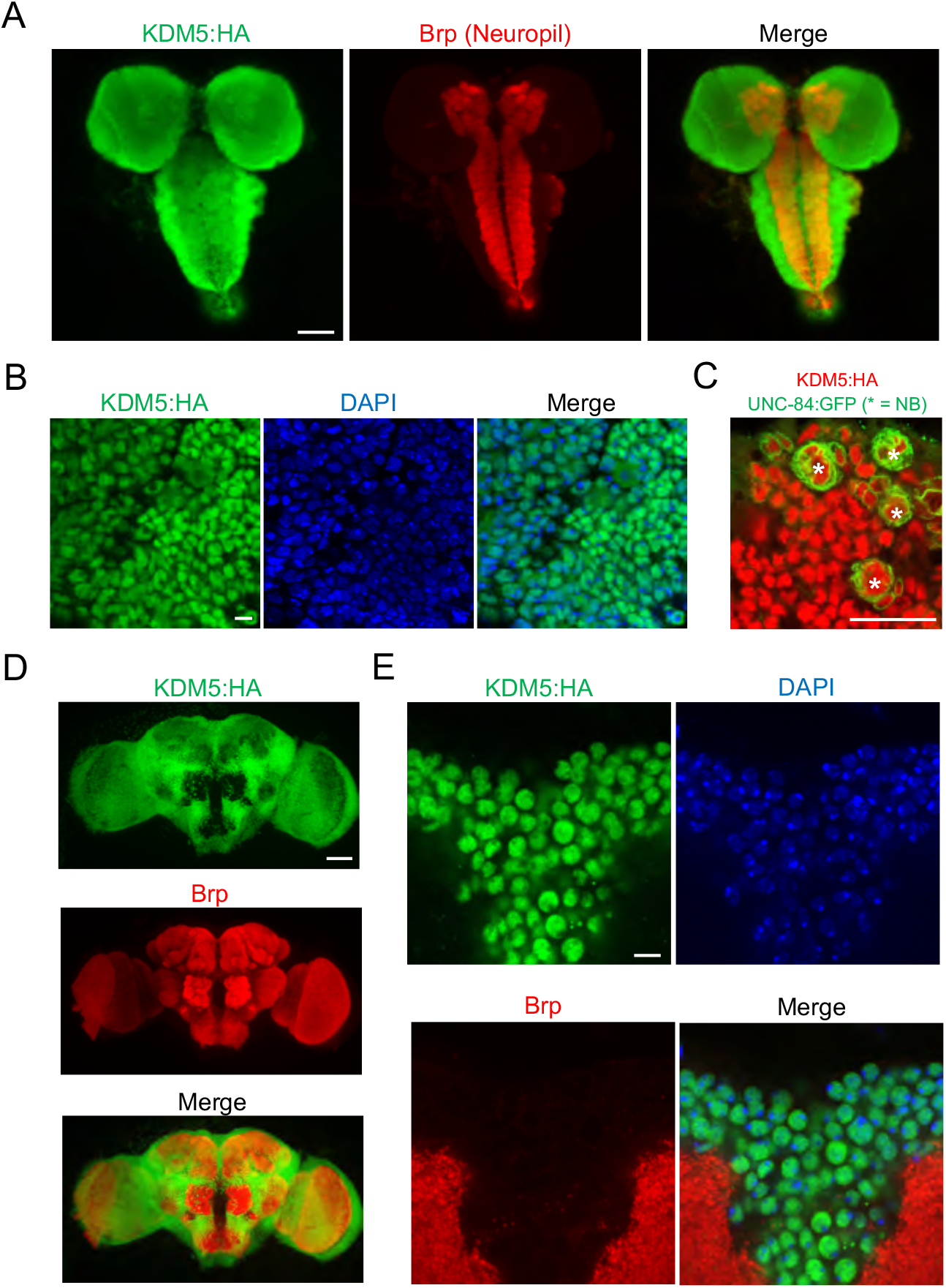
KDM5 is Broadly Expressed in Nuclei of the *Drosophila* Larval CNS and Adult Brain. (A) Maximal Z projection of third-instar larval CNS revealing broad expression of endogenously tagged KDM5:HA and stained with anti-HA and anti-Brp. (B) Cortical region of a larval (WL3) brain lobe stained with anti-HA and DAPI, showing nuclear localization of endogenously tagged KDM5:HA. (C) NB-specific *wor-Gal4* driving expression of UNC-84:GFP to demonstrate endogenously-tagged KDM5:HA expression within nuclei of WL3 central brain NBs (marked by *) and presumptive GMCs. UNC-84:GFP perdures for approximately 2-3 cell divisions. (D) Maximal Z projection of an adult brain revealing broad expression of endogenously tagged KDM5:HA with anti-HA and counterstained for neuropil with anti-Brp. (E) Dorsoanterior cortical region of an adult brain with endogenously tagged KDM5:HA. The nuclear localization of KDM5:HA is revealed by anti-HA, DAPI, and anti-Brp staining. Scale bars represent 50 μm in (A) and (D), 5 μM in (B) and (E), and 20 μM in and (F).

### KDM5 is required within neural precursors and immature neurons for proper MB morphology and behavior

To define the functional requirements of KDM5 during MB development, we utilized an inducible *kdm5* shRNA transgene that we and others have shown effectively reduces KDM5 levels (Chen et al., 2019; Liu et al., 2014; Navarro-Costa et al., 2016). We first knocked down *kdm5* broadly within all MBNBs, MB-GMCs, and Kenyon cells throughout development using *OK107-Gal4* (Fig. 3A,B). This resulted in significant neuromorphological defects of the α/β MB lobes, with the major phenotype being an overextension of the β lobes across the midline (Fig. 3C,D). We next assessed the consequences of *kdm5* depletion within distinct and overlapping subsets of mature Kenyon cells. We hypothesized that if KDM5 functioned within mature, post-mitotic Kenyon cells to maintain MB morphology, then its depletion would lead to neuronal abnormalities. Fas2 staining of animals expressing the *kdm5* shRNA transgene using the mature Kenyon cell-specific Gal4 drivers *C708a-*, *c305a-*, *H24-*, *201Y-*, and *c739-Gal4*, failed to produce any significant gross morphological defects of the α/β lobes (Fig. 3D). KDM5 is therefore unlikely to be required exclusively within mature, post-mitotic Kenyon cells for proper α/β lobe development. Given that concomitant depletion of KDM5 within MBNBs, MB-GMCs and Kenyon cells, but not within mature Kenyon cells, resulted in α/β lobe abnormalities, we suspected that KDM5 may function within NPCs to guide MB development. Knockdown of *kdm5* using two independent NB-restricted *Gal4* driver lines, *worniu-Gal4* (*wor-Gal4*) and *inscuteable-Gal4* (*insc-Gal4*), resulted in profound α/β lobe defects (Fig. 3C,D). Notably, a significant proportion of these brains simultaneously displayed both growth and guidance defects of the α/β lobes. Although *wor-Gal4* expression is NB specific, KDM5 depletion and GFP expression continued to be observed in presumptive GMCs and post-mitotic cells surrounding each NB (Fig. 3E). This is likely attributed to perdurance of the Gal4 activator protein and/or the *kdm5* shRNA. KDM5 could therefore be required within the NB, GMC or even post-mitotically within the immature neuron for proper α/β Kenyon cell neurodevelopment.

**Figure 3.**
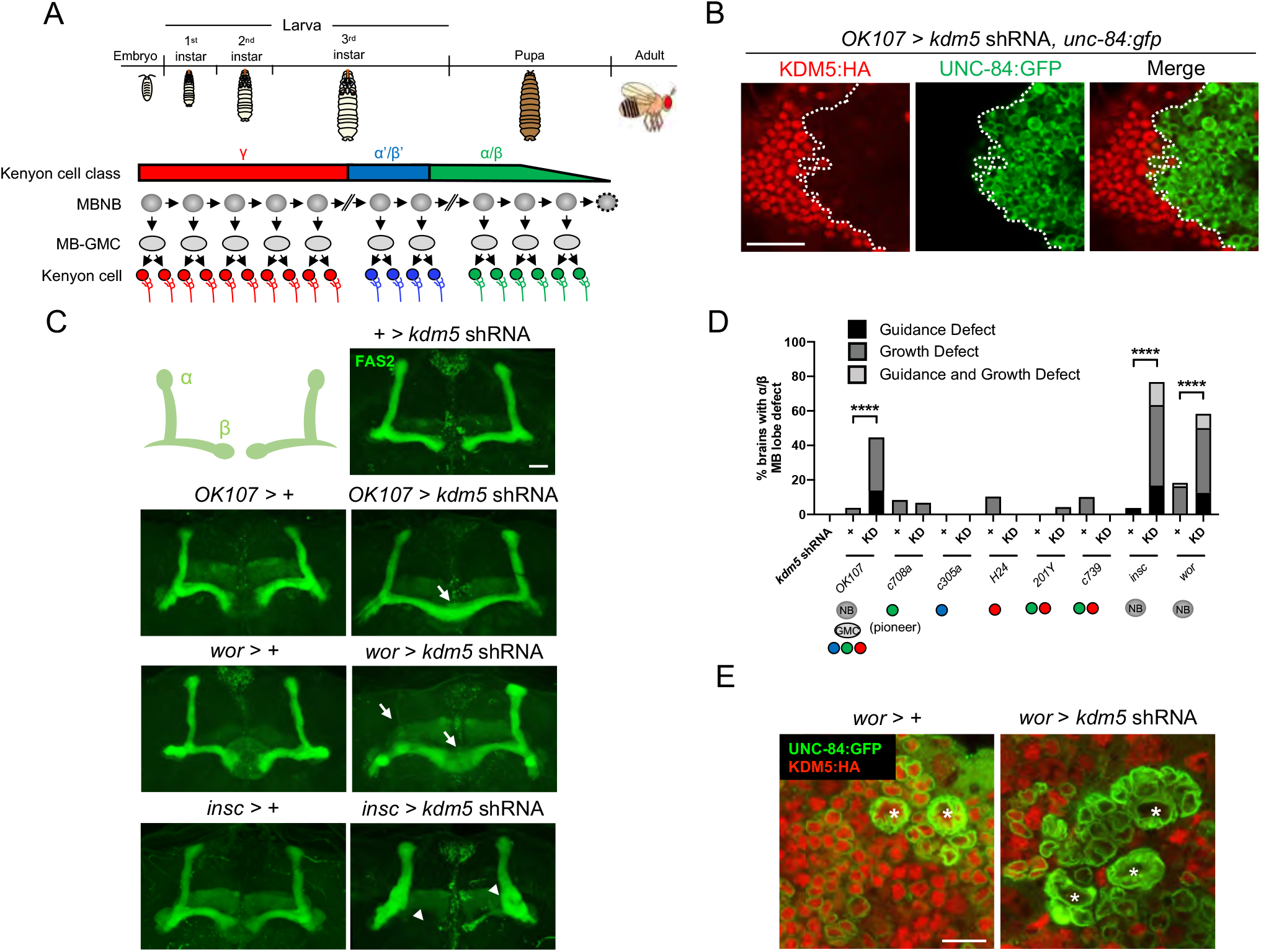
Depletion of KDM5 within Neural Precursors Results in Neuroanatomical Defects of the MB. (A) Schematic showing the sequential generation of distinct subclasses of Kenyon cells throughout development. MBNBs are eliminated *via* apoptosis (dotted line) immediately prior to eclosion. (B) *OK107-Gal4* driving expression of UNC-84:GFP to reveal shRNA-mediated KDM5:HA depletion within adult Kenyon cell nuclei. (C) Representative Z projections of adult *kdm5* knockdown animals exhibiting significant α/β lobe defects and their respective *kdm5* shRNA and *GAL4* controls. The antibody anti-Fas2 is used to visualize α/β lobes. Arrows indicate growth defects and arrowheads indicate guidance defects. (D) Quantification of α/β MB lobe defects in flies expressing *kdm5* shRNA driven by neural progenitor cell- and Kenyon cell-specific drivers. “KD” = shRNA mediated knockdown of *kdm5*. *n* = 16-49 (mean *n* = 29). **** *P* < 0.0001 (chi-square test with Bonferroni Correction). (E) Z projection of larval cortex revealing *wor-Gal4* driven expression of *kdm5* shRNA and *unc-84:gfp transgenes*. KDM5:HA depletion is observed in presumptive GMCs and post-mitotic cells surrounding NBs (marked by an asterisk). Scale bars represent 20 μm in (B) and (C) and 10 μm in (E).

To determine if KDM5 is indeed required within NBs for proper MB development, we used *R71C09-Gal4*, a *Gal4* line which drives expression within GMCs and early born neurons of the *Drosophila* larval CNS (Fig. 4A) (Li et al., 2014; Marshall & Brand, 2017; Aughey et al., 2018). This driver did not appear to drive expression within the majority of NBs observed, including within the MBNBs (Fig. 4A). When *kdm5* was knocked down using *R71C09-Gal4,* KDM5 was dramatically depleted within presumptive GMCs and immature neurons (Fig. 4B). We also noted that *R71C09-Gal4* strongly drove expression in newly born α/β Kenyon cells of recently eclosed adults whose axons lie within the core fibers *of* the MB pedunculus, which was consistent with its strong expression in immature neurons (Fig. 4C). Depletion of *kdm5* using this driver resulted in profound α/β lobe defects, indicating that KDM5 is functionally required within GMCs and immature MB neurons for proper axonal development (Fig. 4D). As knockdown using *R71C09-Gal4* resulted in MB abnormalities, we next used this driver to re-express *kdm5* in GMCs and immature neurons of *kdm5*^*140*^ animals. Re-expression of *kdm5* using *R71C09-Gal4* significantly rescued the defects we had previously observed in *kdm5*^*140*^ animals, further demonstrating that KDM5 acts within GMCs and immature neurons to promote proper MB development (Fig. 4E).

**Figure 4.**
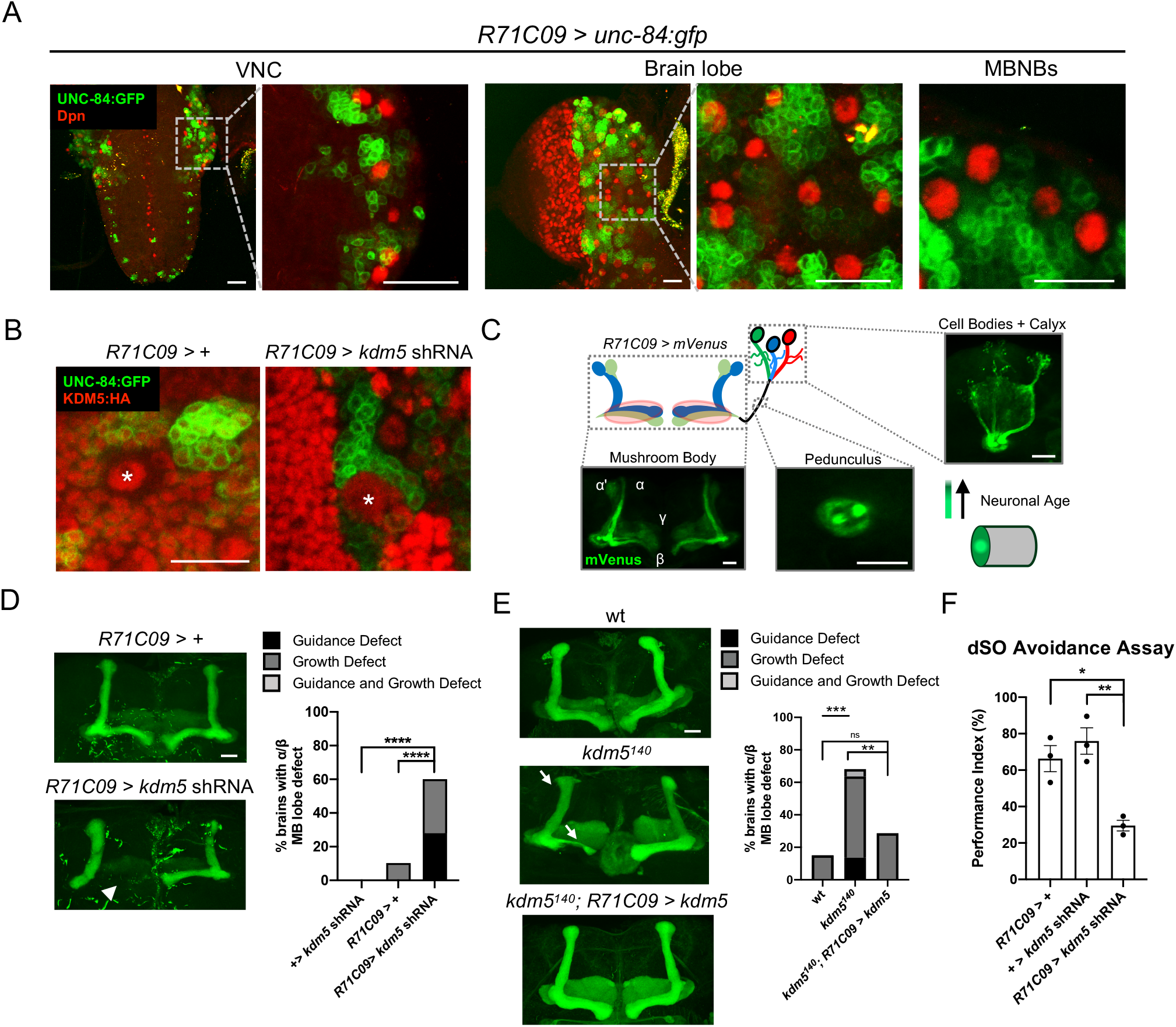
Functional Analysis and Expression Pattern of *R71C09-Gal4* within GMCs and Immature Neurons of the *Drosophila* CNS. (A) Whole mount Z projections of a larval ventral nerve cord (left), brain lobe (middle), and brain cortical region (right). Z projections show *R71C09-Gal4* driven expression of UNC-84:GFP, counterstained with NB-specific anti-Dpn. (B) Z projection of larval cortex revealing *R71C09-Gal4* driven *expression of* UNC-84:GFP with and without *kdm5* shRNA. NBs are marked by an asterisk. (C) Optical sections of an adult MB with its associated pedunculus, Kenyon cell bodies and calyces expressing an *R71C09-Gal4* driven mVenus reporter. *R71C09-Gal4* strongly drives mVenus expression in newly born neurons located within core fibers of the pedunculus. (D) Representative Z projections (left) and quantification (right) of adult α/β MB lobe defects in flies expressing *kdm5* shRNA driven by *R71C09-Gal4*. The antibody anti-Fas2 is used to visualize α/β lobes. *n* = 15-39 (mean *n* = 26). Arrowhead indicates a guidance defect. **** *P* < 0.0001 (chi-square test with Bonferroni Correction). (E) Representative α/β lobe Z projections and quantification of pharate wildtype (*NP4707*^*rev2*^ revertant), *kdm5*^*140*^, and *kdm5^140^; R71C09 > kdm5* rescue strains. Arrows indicate growth defects and arrowheads indicate guidance defects. The α/β lobes are revealed with anti-Fas2. *n* = 20-28 (mean *n* = 24). ** *P* < 0.01; *** *P* < 0.001 (chi-square test with Bonferroni Correction). (F) Performance Index quantifying dSO avoidance in flies expressing *kdm5* shRNA driven by *R71C09-Gal4*. Emitted dSO is from 100 agitated wild-type flies. Bar graph represents the mean ± SEM for three trials per genotype with responder *n* = 47-57. \**P* < 0.05; \*\**P* < 0.01 (one-way ANOVA with Tukey’s post-hoc test). Scale bars represent 20 μm.

To assess whether these MB defects were associated with behavioral deficits, we utilized a robust two-choice behavioral assay which measures avoidance to *Drosophila* stress odorant (dSO), an innately repulsive sensory cue produced by conspecifics upon mechanical agitation (Fernandez et al., 2014; Suh et al., 2004). Assays quantifying dSO avoidance have routinely been used to measure complex behavioral outputs such as sensory processing and social avoidance (Androschuk et al., 2018; Chen et al., 2019; Fernandez, 2017; Fernandez et al., 2014; Wise et al., 2015). For example, depletion of *fmr1* within the MB has been shown to impair dSO avoidance (Androschuk et al., 2018; Wise et al., 2015). Additionally, ubiquitous depletion of KDM5 using a transheterozygous combination of hypomorphic *kdm5* alleles (*kdm5*^*K6801/10424*^) has been shown to result in deficits to dSO avoidance (Chen et al., 2019). We found that *R71C09-Gal4*-mediated knockdown of *kdm5* resulted in a significant decrease in dSO response compared to that of controls (Fig. 4F). Loss of KDM5, specifically in GMCs and immature neurons, therefore impacts behavior, possibly *via* alterations to MB morphology.

### Transcriptional profiling using Targeted DamID (TaDa) analyses reveals KDM5-regulatory networks critical for neurodevelopment

To define the transcriptional programs regulated by KDM5 within GMCs and immature neurons to regulate MB development we utilized a technique known as Targeted DamID (TaDa), which allows for *in vivo* transcriptional profiling in a cell-type-specific and temporally-controlled manner (Marshall et al., 2016; Southall et al., 2013). By heterologously expressing *E. coli*-derived DNA adenine methyltransferase (Dam) fused to *Drosophila* RNA Polymerase II (Pol II) in MB-GMCs and immature Kenyon cells *via* the Gal4/UAS system, we surveyed gene expression changes resulting from loss of KDM5. Dam-Pol II methylates adenine residues (m6A) at GATC motifs in close proximity to Pol II-occupied DNA, providing a surrogate for actively transcribed loci when normalized to expression of Dam alone.

We expressed Dam-Pol II or Dam in GMCs and immature neurons of late third-instar larvae and pupae of homozygous *kdm5*^*140*^ animals using the *R71C09-Gal4* driver (Fig. 5A). We rationalized that induction of the TaDa system during this developmental window would predominantly result in the transcriptomic profiling of MB-GMCs and immature α/β Kenyon cells, as this is one of the few cell types undergoing extensive neurogenesis during this time period and expresses *R71C09-Gal4* (Heisenberg and Technau, 1982; Ito and Hotta, 1992; Truman and Bate, 1988). Importantly, the temperature shifts required to induce the TaDa system in a *kdm5*^*140*^ animals did not alter the frequency or type of structural MB defects observed (Fig. S1A,B).

**Figure 5.**
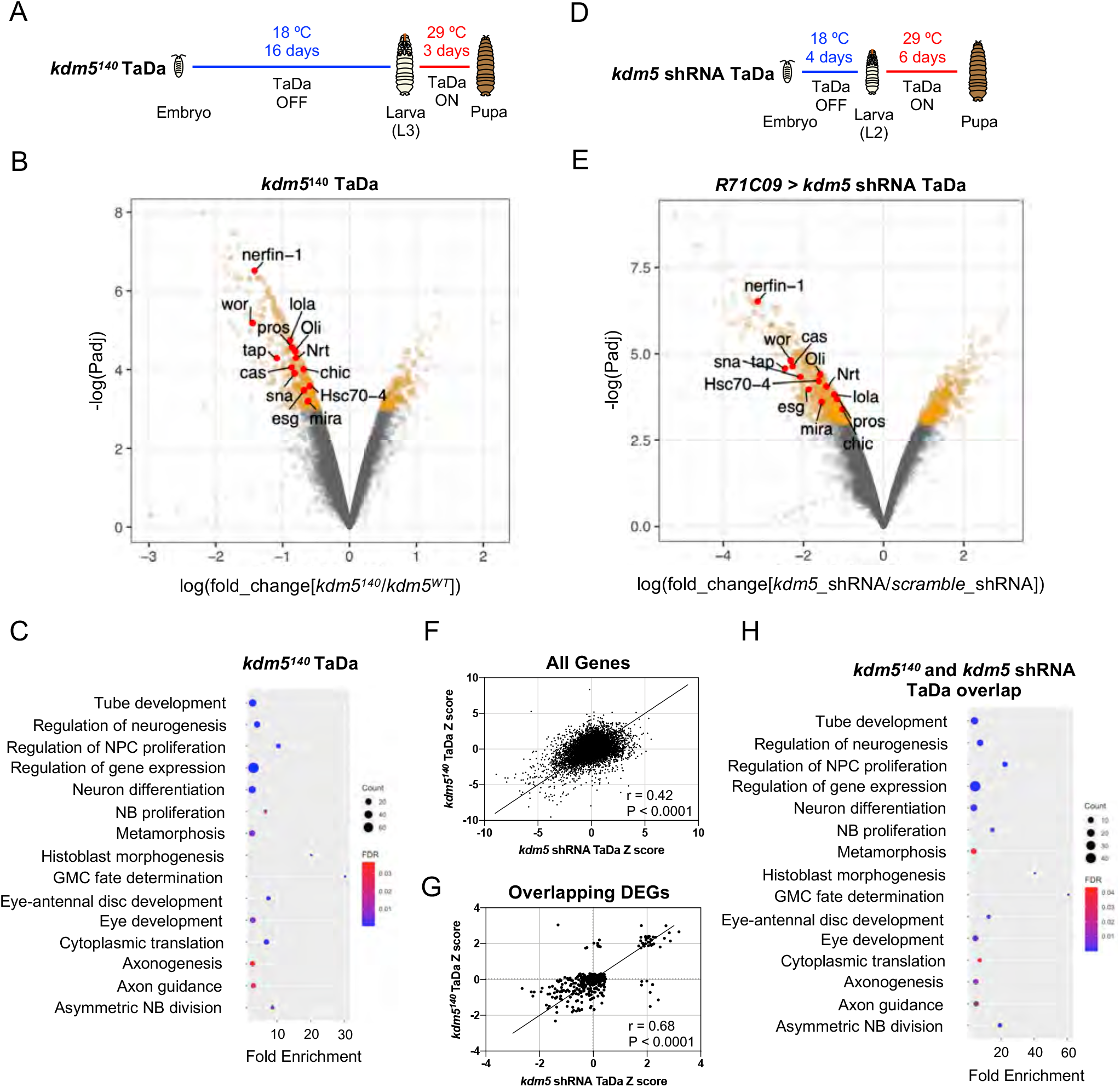
Transcriptome Profiling of KDM5-Depleted GMCs and Immature Neurons by Targeted DamID (TaDa) Reveals KDM5-Regulatory Networks Critical for GMC Proliferation and Neurodevelopment. (A) Timeline of TaDa induction within GMCs and immature neurons of *kdm5*^*140*^ WL3 animals and pupae. (B) Volcano plot of DEGs within GMCs and immature neurons of *kdm5*^*140*^ animals compared to wildtype. Genes with an FDR < 0.05 are in red, with those labeled involved in GMC proliferation and neurodevelopment. TaDa analyses were performed in quintuplicate. (C) Representative distribution of ontology terms for DEGs in *kdm5*^*140*^ GMCs and immature neurons using a PANTHER Overrepresentation Test (Fisher’s exact test with FDR < 0.05). (D) Timeline of TaDa and *kdm5* shRNA induction within GMCs and immature neurons of WL3 animals and pharate adults. (E) Volcano plot of DEGs within *kdm5* shRNA GMCs and immature neurons compared to those expressing a scrambled shRNA. Genes with an FDR < 0.05 are in red, with those labeled involved in GMC proliferation and neurodevelopment. TaDa analyses were performed in triplicate. (F) Correlation of Z scores between DEGs of *kdm5*^*140*^ and *kdm5* shRNA TaDa datasets (Deming regression; *P* < 0.0001). (G) Correlation of Z scores between overlapping DEGs of *kdm5*^*140*^ and *kdm5* shRNA TaDa datasets (Deming regression; *P* < 0.0001). (H) Representative distribution of ontology terms for DEGs in overlapping *kdm5*^*140*^ and *kdm5* shRNA TaDa datasets utilizing a PANTHER Overrepresentation Test (Fisher’s exact test with FDR < 0.05).

Carrying out TaDa from five biological replicates, we identified a total of 636 differentially expressed genes (DEGs) using a statistical cutoff of FDR < 0.05, 438 of which were downregulated and 198 of which were upregulated (Fig. 5B, Table S1). Consistent with previously published mRNA-seq data in *Drosophila* and other organisms, loss of KDM5 resulted in moderate changes to gene expression, with an average 2.4-fold decrease among downregulated genes and 2.1-fold increase among upregulated genes (Drelon et al., 2018; Iwase, Brookes, and Agarwal et al., 2016; Liu et al., 2014; Liu & Secombe, 2015; Lloret-Llinares et al., 2012; Lopez-Bigas et al., 2008; Lussi et al., 2016; Zamurrad et al., 2018). Thus, KDM5 likely functions within this cellular subpopulation to fine-tune gene expression across a number of pathways that are critical for MB development.

To determine if these dysregulated genes were enriched for functional categories, we used the program GeneOntology (The Gene Ontology Consortium, 2000, 2019) which utilizes the PANTHER Classification System (Mi et al., 2013) to mine gene ontology (GO) information. This revealed significant enrichment for categories involved in GMC fate determination (e.g. *pros, brat, mira, cas*), neural precursor cell proliferation (e.g. *pros, wor, ase, esg, sna, E(spl)mγ-HLH, E(spl)mβ-HLH*), axon guidance (e.g. *pros, dac, brat, Hr51, chic, tap, pdm3, nerfin-1, lola*), and cytoplasmic translation, among others (Fig. 5C; Table S2). The finding that ribosomal protein genes, such as *RpS2*, *RpS24*, *RpS28b*, *RpL3*, *Rpl23*, *RpL39*, and *RpL41*, were altered in our *kdm5*^*140*^ TaDa data is consistent with our previously reported RNA-seq data from adult heads of demethylase-dead *kdm5* strains showing that genes required for cytoplasmic translation were affected (Zamurrad et al., 2018). Interestingly, however, knockdown of the transcription factor *myc*, which regulates the expression of most ribosomal protein genes (Bellosta and Gallant, 2010; Grewal et al., 2005), in MBNBs, GMCs, and Kenyon cells using the *OK107-Gal4* driver did not result in any gross morphological defects of the α/β lobes (Fig. S2). These data show that our observed MB phenotypes are not due to the regulation of cytoplasmic translation by KDM5.

H3K4me3 marks have been shown to be associated with regions of accessible chromatin, which often contain regulatory DNA sequences such as promoters and enhancers (Bhaumik et al., 2007; Li et al., 2012; Park et al., 2020; Thurman et al., 2012). If KDM5 functions independent of its H3K4me3 demethylase activity within GMCs and immature neurons to regulate MB development, we would hypothesize that loss of KDM5 would have minimal effects on chromatin accessibility within this cellular subpopulation. To measure changes to chromatin landscape upon KDM5 loss, we utilized Chromatin Accessibility profiling using Targeted DamID (CATaDa) (Aughey et al., 2018, 2019). Dam methylates regions of highly accessible chromatin, thus providing an *in vivo* surrogate for chromatin accessibility (Aughey et al., 2018, 2019). Comparing Dam methylation levels in *R71C09-Gal4* expressing cells between *kdm5*^*140*^ and wild-type animals revealed minor changes to chromatin accessibility at a cutoff of FDR < 0.01 (Fig. S3A,B). Overlapping the reduced accessibility regions with the downregulated genes from our TaDa analysis revealed only 17 genes associated with reduced chromatin accessibility in GMCs and immature neurons at a cutoff of FDR < 0.01 (Fig. S3C). Although this represented a significantly enriched proportion of the total number genes associated with changes in chromatin accessibility (*P* = 4.787E-09), they were not enriched *via* GO analysis for any biological categories. Thus, it is likely that KDM5 functions largely *via* demethylase independent transcriptional mechanisms to regulate MB morphology without dramatically altering chromatin accessibility.

Since *kdm5*^*140*^ animals have chronic loss of *kdm5* in all tissues, the gene expression changes we observed may be the result of both cell-autonomous and non-cell-autonomous effects. To look directly at the KDM5-regulated transcriptome in GMCs and early born neurons in a cell-type specific manner, we performed TaDa in animals expressing a *kdm5* shRNA transgene under the control of *R71C09-Gal4*. In addition to utilizing a Dam-only control, we also accounted for activation of the RNAi pathway by expressing a scramble shRNA transgene under identical conditions. To ensure that changes to gene expression were reflective of the severe MB phenotypes we had previously observed, we induced the expression of *kdm5* shRNA and *dam:pol ll* for six days, beginning during the early second-instar larval (L2) stage, to allow for sufficient depletion of KDM5 within MB-GMCs and immature α/β Kenyon cells (Fig. 5D). This temporally-targeted knockdown strategy recapitulated the adult α/β lobe phenotypes we had previously observed for constitutive *kdm5* knockdown with *R71C09-Gal4* (Fig. S4A,B). From three biological replicates and a statistical cutoff of FDR < 0.05, this TaDa experiment identified 1069 DEGs, 659 of which were downregulated and 410 of which were upregulated (Fig. 5E, Table S3). Compared to the *kdm5*^*140*^ TaDa, the greater number of DEGs from the *kdm5* RNAi TaDa could be attributed to the extended duration of the *kdm5* knockdown and induction of Dam:Pol II. Additionally, we observed larger changes to gene expression, with an average 5.5-fold decrease among downregulated genes and a 4.3-fold increase among upregulated genes (Fig. 5E, Table S3). This could be ascribed to the acute loss of *kdm5* expression due to RNAi-mediated depletion which would decrease the likelihood of compensatory changes occurring.

To obtain a list of high-confidence genes regulated by KDM5, we compared the DEGs found in our *kdm5*^*140*^ and *kdm5* knockdown TaDa datasets. This revealed a total of 335 overlapping dysregulated genes, 319 of which were up- or downregulated in both datasets (r=0.68, P < 0.0001) (Fig. 5F,G). GO analysis of the 319 similarly dysregulated DEGs revealed an even greater enrichment in categories related to GMC fate determination (*pros, mira, cas*), neural precursor cell proliferation (*wor, esg, pros, mira, sna*), and axon guidance (*pros, chic, lola, Oli, tap, Nrt, nerfin-1, Hsc70-4*), among others (Fig. 5H; Table S4). Interestingly, only 17 of these genes overlapped with our previously published *kdm5*^*140*^ wing disc RNA-seq data (Drelon et al., 2018) (Fig. S5A,B; Table S5). This nonsignificant overlap (Pearson’s r = 0.09) suggests that KDM5 functions in a predominantly tissue- and cell-specific manner to regulate the expression of downstream targets.

To identify biologically relevant functional networks for the 319 KDM5-regulated, high confidence genes, we performed gene network and community clustering analyses (Morris et al., 2011; Shannon et al., 2003). These analyses revealed seven discrete networks with greater than two nodes (Fig. 6). Of these, two networks were highly enriched for genes implicated in MB development (*toy* and *tap*), neural precursor cell proliferation (*pros, ase, wor, mira, sna, esg, E(spl)mγ-HLH,* and *E(spl)mβ-HLH*), and axon growth and guidance (*pros, nerfin-1, tap, emc, Nrt,* and *elav*). Notably, one gene, *prospero* (*pros*), encodes a homeodomain-domain containing transcription factor that is a critical regulator of axon pathfinding and growth in other neuronal contexts, and belonged to multiple ontology categories that were relevant to our analyses (Doe et al., 1991; Froldi et al., 2015; Tea et al., 2010; Vaessin et al., 1991).

**Figure 6.**
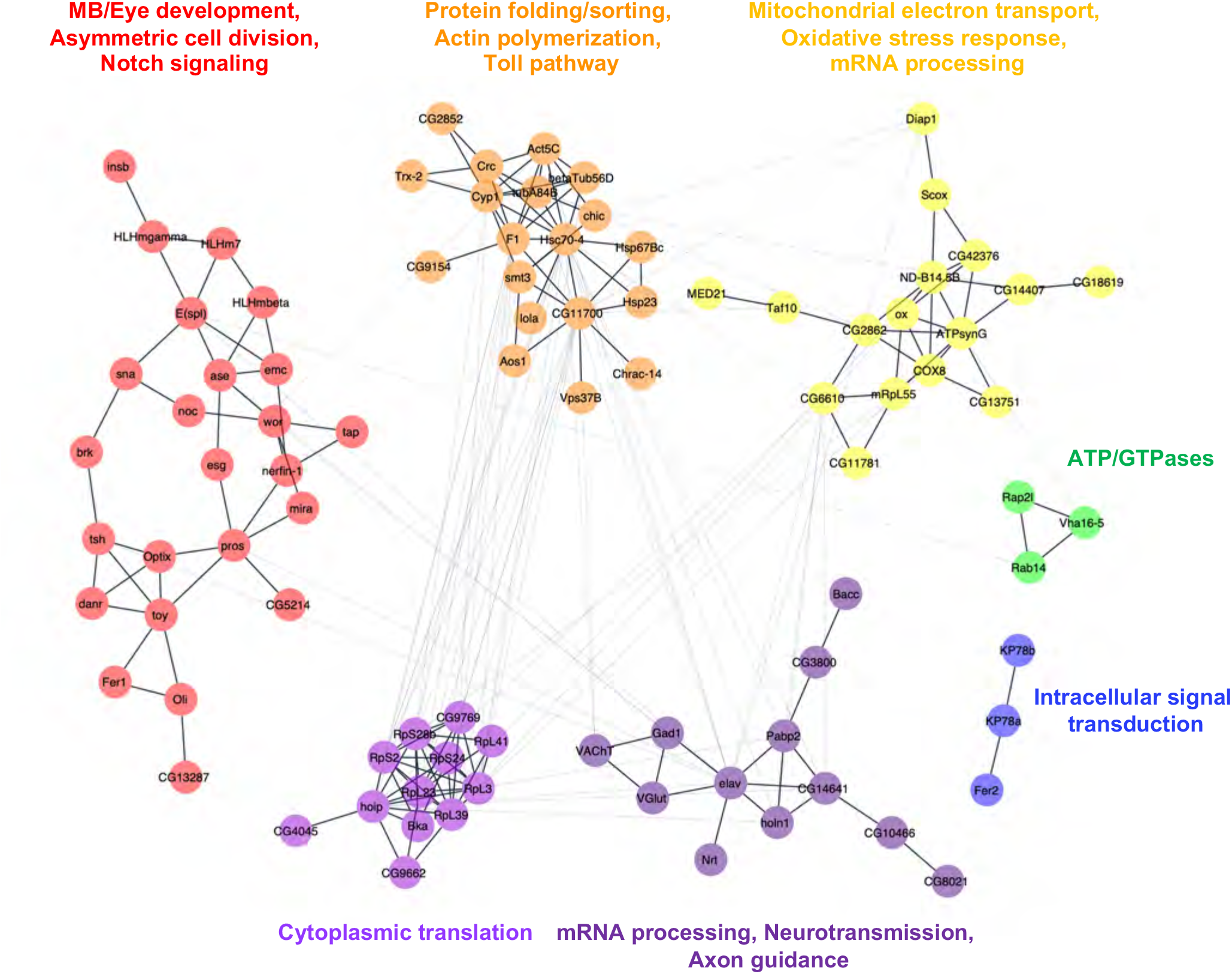
Network of Known and Predicted interactions between Overlapping DEGs of *kdm5*^*140*^ and *kdm5* shRNA TaDa Datasets. Gene network analysis and community clustering were performed using Cytoscape with a minimum confidence score of 0.4. Networks with greater than two nodes are shown and color-coded based on cluster. Labels indicate general categories of overlapping *kdm5*^*140*^ and *kdm5* shRNA TaDa DEGs within each cluster.

### The transcription factor *pros* genetically interacts with *kdm5* and is required for proper MB development

Our gene expression data pointed to a link between KDM5 and Pros in the regulation of MB development. We therefore used a recently published TaDa dataset of direct Dam:Pros targets within GMCs and immature neurons using *R71C09-Gal4* to look for an enrichment of Pros-bound targets within KDM5-regulated genes (Liu et al., 2020; GEO accession number GSE136413). Strikingly, 45% (142 of 319) of KDM5-regulated genes were bound by Pros (*P* = 2.43E-64, Fisher’s exact test) (Fig. 7A; Table S6). These data suggest that Pros could be a key mediator of KDM5 function in GMCs and immature neurons. Our data also implicate Pros in the activation of genes, as the majority of Pros-bound genes were downregulated upon KDM5 depletion (Fig. 7B).

**Figure 7.**
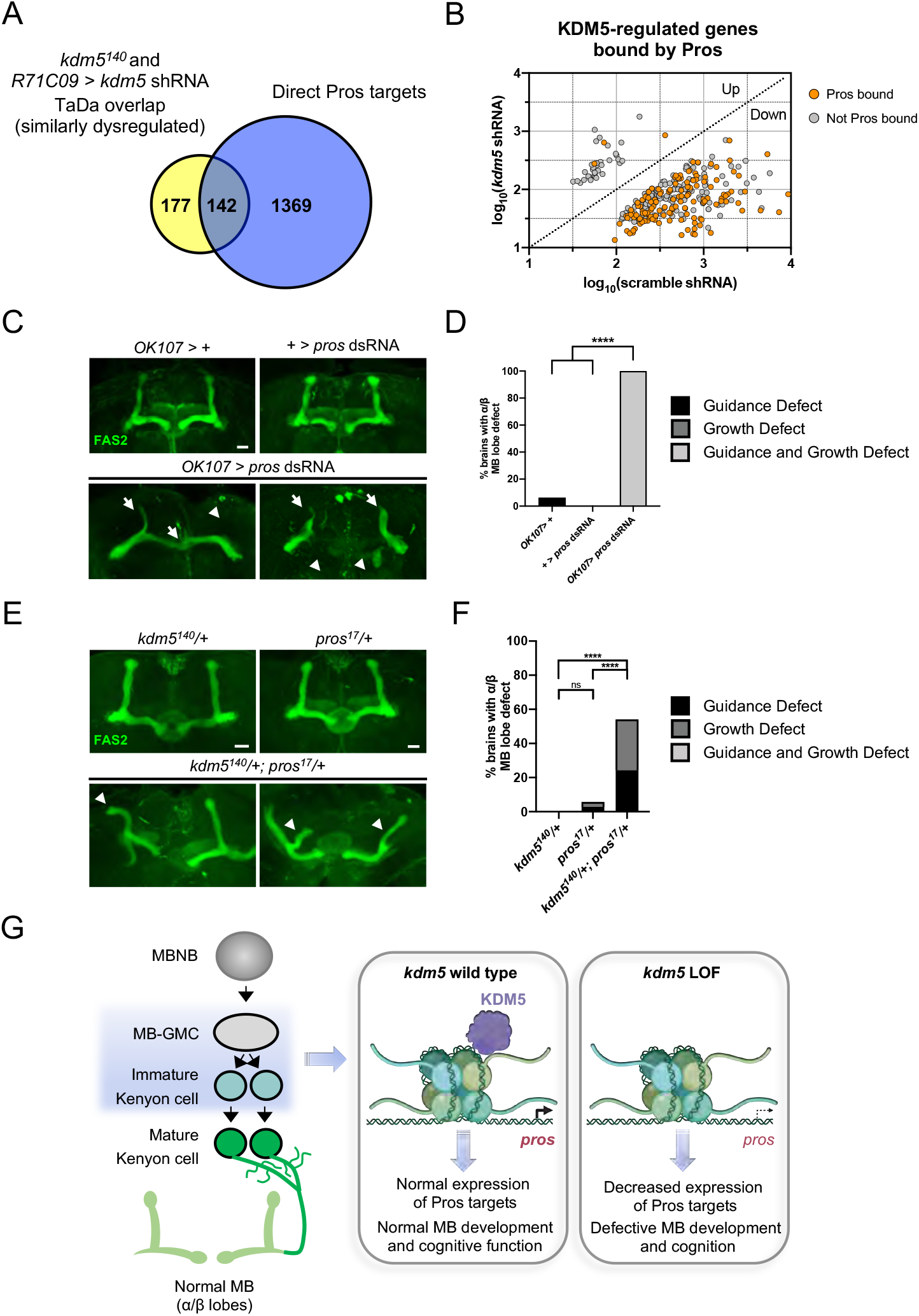
Neuromorphological and Transcriptomic Analyses Reveal a Genetic Interaction between Prospero and KDM5. (A) Venn diagram illustrating intersection of similarly-dysregulated *kdm5*^*140*^ and *kdm5* shRNA overlapping DEGs with a previously published Dam:Pros TaDa binding dataset (Fisher’s exact test, *P* = 2.43E-64) (Liu et al., 2020). (B) Analysis of the 319 similarly-dysregulated *kdm5*^*140*^ and *kdm5* shRNA overlapping DEGs (with values plotted from the *kdm5* shRNA TaDa). Direct Pros targets from the previously published Dam:Pros TaDa dataset (Liu et al., 2020) are labeled in orange. (C) Representative Z projections of *OK107* > *pros* RNAi adults exhibiting significant α/β lobe defects. The α/β lobes are revealed by anti-Fas2. (D) Quantification of α/β MB lobe defects in flies expressing *pros* shRNA driven by *OK107-Gal4*. *n* = 16-19 (mean *n* = 17). **** *P* < 0.0001 (chi-square test with Bonferroni Correction). (E) Representative Z projections of representative *kdm5*^*140*^/+ and *pros*^*17*^/+ heterozygous adult α/β MB lobes (top) and *kdm5*^*140*^/+; *pros*^*17*^/+ transheterozygous adult α/β MB lobes (bottom). The α/β lobes are revealed by anti-Fas2. (F) Quantification of α/β MB lobe defects in *kdm5*^*140*^/+; *pros*^*17*^/+ transheterozygous adults and heterozygous controls. *n* = 28-37 (mean *n* = 34). **** *P* < 0.0001 (chi-square test with Bonferroni Correction). (G) Model proposing a genetic interaction between *pros* and *kdm5* within GMCs and immature α/β Kenyon cells. Loss of *kdm5* leads to downregulation of *pros* and its targets, resulting in defects to MB neurodevelopment and cognitive function. Image was created with BioRender. Scale bars represent 20 μm.

While roles for Pros in regulating NB and GMC asymmetric division are well-described (Choksi et al., 2006; Doe et al., 1991; Vaessin et al., 1991), its role in MB development remains unexplored. We thus tested the effects of Pros depletion in the MB with the expectation that it would phenocopy the MB defects we had observed in *kdm5*^*140*^ pharate and *kdm5* shRNA-expressing adult brains. *R71C09-Gal4*-mediated knockdown of Pros resulted in early larval lethality, preventing us from examining MB phenotypes. However, depleting Pros within the smaller cellular population of MB progenitor and Kenyon cells using *OK107-Gal4* allowed for the formation of viable adults. Analyses of these adult brains revealed a complete disruption of MB neurodevelopment, with both growth and guidance defects present in 100% of observed MBs (Fig. 7C,D).

As *kdm5* or *pros* depletion within MB progenitor and Kenyon cells each independently result in significant axonal defects, we next assessed whether *kdm5* and *pros* functioned within the same genetic pathway. To test this, we generated animals heterozygous for null alleles of both *kdm5* and *pros*, thereby simultaneously reducing levels of both proteins. Animals heterozygous for either *kdm5*^*140*^ or *pros*^*17*^ did not display significant MB defects. In contrast, adults that were transheterozygous for *kdm5*^*140*^ and *pros*^*17*^ displayed MB abnormalities, albeit less severe than those of *kdm5*^*140*^ homozygous pharates or OK107-Gal4-mediated *pros* knockdown adults alone (Fig 7E,F). These data are consistent with KDM5 and Pros functioning synergistically to affect MB development. KDM5 is therefore likely to function within GMCs and immature neurons to regulate levels of *pros* and its downstream targets, thereby driving MB development and promoting proper cognitive function (Fig. 7G).

## DISCUSSION

Here, we utilize an *in vivo* transcriptional approach coupled with neuromorphological and behavioral analyses to demonstrate that KDM5 is essential for the development and function of the adult MB. Although previous work in murine models has shown that knockout of the *kdm5* ortholog *Kdm5c* leads to dendritic spine abnormalities in cortical brain regions, these studies utilized ubiquitous knockout systems which were not amenable to spatiotemporal genetic manipulation (Iwase, Brookes, and Agarwal et al., 2016; Vallianatos et al., 2020). *Drosophila*, however, provide a genetically tractable system to modulate gene expression in a spatiotemporally restrictive manner. Using the MB as a model for neuronal development, we demonstrate that KDM5 expression within GMCs and immature neurons is necessary for proper neuropil development. Depletion of KDM5 within mature subpopulations of Kenyon cells did not affect gross MB structure, suggesting that there exists a critical neurodevelopmental window during which KDM5 is required to regulate genetic programs essential for axonal growth and guidance. Notably, the MB growth and guidance defects we observed phenocopied those seen in other *Drosophila* ID models, such as the *fmr1* and *dnab2* RNA binding protein mutants (Kelly et al., 2016; Michel, 2004). It is thus possible that ID genes may function synergistically within common neurodevelopmental pathways to affect neuronal architecture and function – a hypothesis that is currently under active investigation (van Bokhoven, 2011; Fenckova et al., 2019; Kochinke et al., 2016; Najmabadi et al., 2011).

We additionally performed TaDa analyses to survey gene expression changes resulting from loss of *kdm5* in a cell-specific and temporally-restricted manner. Although previous transcriptomic studies from us and others have utilized extracted RNA from dissected tissues or from neuronal or fibroblast cultures as input for RNA-seq (Brookes et al., 2018; Iwase, Brookes, Agarwal, Badeaux, et al., 2016; Liu & Secombe, 2015; Scandaglia et al., 2017; Vallianatos et al., 2020; Wei et al., 2016; Zamurrad et al., 2018), this is the first study describing changes to gene expression resulting from cell-specific loss of *kdm5*. This technique allowed us to survey gene expression changes within GMCs and immature neurons of *kdm5* null and cell-type specific knockdown animals, revealing a number of differentially expressed genes associated with neurodevelopment. Although prior work from *C. elegans* led to the postulate that KDM5 regulates neuronal development by repressing expression of the actin regulator *wsp-1*/*WASp* (Mariani et al., 2016), our TaDa analyses did not find altered expression of the *Drosophila wsp-1* ortholog or its family members. Interestingly, CATaDa analyses revealed only minimal changes to chromatin accessibility within GMCs and immature neurons upon *kdm5* loss. Taken together, our data suggest that KDM5 may regulate the development of different neuronal cell-types by modulating the expression of distinct subsets of genes in a demethylase independent manner.

Our data revealed a genetic link between KDM5 and the homeodomain-containing transcription factor Pros. Pros is a cell fate determinant that is expressed in most neuronal precursors and immature neurons and is involved in cell-cycle exit, neuronal differentiation and axonal development (Choksi et al., 2006; Doe et al., 1991; Froldi et al., 2015; Tea et al., 2010; Vaessin et al., 1991). Loss of *pros* results in axonal routing defects within embryonic motor and sensory neurons (Vaessin et al., 1991) and can lead to the miswiring of olfactory projection neuron dendrites (Tea et al., 2010). Despite its demonstrated importance in other neuronal contexts, a role for Pros in regulating MB growth and guidance programs was previously uncharacterized. We demonstrate that depletion of *pros* within MBNBs, MB-GMCs, and immature Kenyon cells, leads to severe MB growth and guidance defects. Importantly, Pros is expressed primarily in neural precursors and immature neurons at a time when KDM5 function is critical for MB development (Doe et al., 1991; Vaessin et al., 1991).

Our transcriptome data additionally revealed the dysregulation of a number of Pros interactors upon *kdm5* depletion. For example, the direct Pros target *nerfin-1*, which encodes a zinc finger transcription factor required for early axon pathfinding by most CNS neurons in the *Drosophila* embryo (Kuzin et al., 2005; Liu et al., 2020), was significantly downregulated. Additionally, genes encoding Toy and Tap, which have been shown to regulate MB morphology (Furukubo-Ttokunaga et al., 2009; Yuan et al., 2016), were similarly affected. Transcriptional dysregulation of *pros* and its targets may thus provide important molecular insight into the neuronal and cognitive phenotypes associated with *kdm5* loss of function.

Although we demonstrate a genetic link between *kdm5* and *pros* in regulating MB development, the exact mechanism(s) through which KDM5 accomplishes this remains unknown. We predict, however, that the regulation of *pros* by KDM5 is independent of its canonical histone demethylase activity, as animals lacking this enzymatic function have MBs that are phenotypically indistinguishable from those of wildtype (Zamurrad et al., 2018). Flies lacking KDM5 histone demethylase activity do, however, have cognitive deficits (Zamurrad et al., 2018), suggesting that KDM5 may regulate neuronal development and function *via* multiple mechanisms. Furthermore, our CATaDa analysis of GMCs and immature neurons did not reveal significant changes to chromatin accessibility that were enriched for any biological processes upon *kdm5* loss. Our data are therefore consistent with a model in which KDM5 engages distinct transcriptional programs in neurons and neural progenitors depending on developmental context.

Consistent with this model, several ID-associated mutations in KDM5C have been shown to alter H3K4me3 levels *in vitro*, whereas others do not (Brookes et al., 2015; Vallianatos et al., 2018). Based on these data and the results reported here, it is likely that ID-causing mutations in human *KDM5A*, *KDM5B* or *KDM5C* might fall into three classes: the first may only affect KDM5 demethylase activity, the second may only affect non-enzymatic activities necessary for transcriptional regulation, and the third may affect both. The extent to which canonical and non-canonical activities are disrupted may, indeed, correlate with the severity of the cognitive deficit or presence of syndromic features. Future work will further elucidate how individual patient mutations may disrupt these transcriptional regulatory mechanisms to influence neuronal and behavioral outputs, thus providing a promising strategy for predicting and treating KDM5-associated ID.

## ACKNOWLEDGMENTS

We thank Nicholas Baker, Hannes Bülow, and all members of the Secombe Lab for their feedback and edits on the manuscript. We additionally thank Hannes Bülow, Andreas Jenny, Bernice Morrow, Anna Francesconi, and Nicholas Baker for their suggestions and feedback during the formulation and execution of the above experiments. We appreciate the confocal microscope training and assistance provided to us by Hillary Guzik and members of the Einstein Analytical Imaging Facility (AIF). Portions of this study would not have been possible without fly strains and reagents generously donated to us by Gilbert Henry, Andrea Brand, Lily Yeh Jan and Yuh Nung Jan. Stocks obtained from the Bloomington Drosophila Stock Center (NIH P40OD018537) were also used in this study. The 1D4 and 9.4A monoclonal antibodies were obtained from the Developmental Studies Hybridoma Bank, created by the NICHD of the NIH and maintained at The University of Iowa. We are additionally grateful to the NIH Special Instrument Grant 1S10OD023591-01 and the Cancer Center Support Grant P30CA013330. This research was supported by the NIH Ruth L. Kirschstein National Research Service Award F31NS110278, the Einstein MSTP Training Grant T32GM007288, and the Junior Investigator in Neuroscience Research Award (JINRA) from the Dominick P. Purpura Department of Neuroscience to H.A.M.H., NIH R01GM112783 to J.S., and NHMRC grants APP1128784 and APP1185220 to O.J.M.

## AUTHOR CONTRIBUTIONS

H.A.M.H. and J.S. conceived of and designed the experiments. H.A.M.H. conducted all experiments and H.M.B. generated the endogenously-tagged *kdm5:HA* strain. H.A.M.H. analyzed the immunohistochemistry and behavioral data and H.A.M.H., O.J.M., and J.S. analyzed the TaDa data. H.A.M.H., O.J.M., and J.S. wrote the paper and secured funding.

## MATERIALS AND METHODS

### Resource Availability

#### Lead Contact

Further information and requests for resources and reagents should be directed to and will be fulfilled by the Lead Contact, Julie Secombe.

#### Materials Availability

The *kdm5:HA* strain generated in this study is available from the Lead Contact without restriction. DamID-Seq data have been deposited in the Gene Expression Omnibus (GEO) under accession number GSE156010.

#### Data and Code Availability

The code supporting the current study is available at http://marshall-lab.org/code/.

### Fly Strains and Genetics

A detailed list of the genotypes of the flies used in each figure is included in the Key Resources Table.

All *GAL4* and *GMR GAL4* lines were generated at the Janelia Research Campus/HHMI (Pfeiffer et al., 2008; Jenett et al., 2012) and were obtained from the Bloomington *Drosophila* Stock Center (BDSC) at Indiana University.

The following transgenes were used: *UAS-kdm5-shRNA* (RRID: BDSC_35706), *20XUAS-IVS-CsChrimson.mVenus* (RRID: BDSC_55136), *UAS-tub-GAL80^ts^* (RRID: BDSC_7108), *UAS-pros-dsRNA* (RRID: BDSC_42538), *UAS-myc-dsRNA* (VDRC stock# KK106066). The *UAS-LT3-dam* and *UAS-LT3-dam-pol II* lines were kindly shared by Andrea Brand (U. Cambridge, Gurdon). The *5XUAS-unc84-2Xgfp* line was kindly shared by Gilbert Henry and Todd Laverty (HHMI Janelia). The *kdm5*^*140*^ mutant allele and *UAS-kdm5* transgene have been previously described (Drelon et al., 2018; Secombe et al., 2007).

### Cloning and Transgenesis

To tag the endogenous *kdm5* locus with three in-frame *HA* epitope tags, we used CRISPR/Cas9-mediated knock-in. The *HA* epitopes and the homology arms, carrying a synonymous mutation for the PAM sequence, were PCR amplified from a clone containing the *kdm5* locus from the wild type strain *w*^*1118*^ (pattB.gkdm5; Navarro-Costa et al. 2016). The donor DNA repair template consisted of three PCR fragments cloned, by In-Fusion^®^ HD (Takara, Bio), into the pHD-ScarlessDsRed vector (Addgene plasmid #51434). AarI and SapI enzymes were used to linearize the plasmid. The flyCRISPR Optimal Target Finder tool was used to select the target genomic cleavage and design the gRNA (http://targetfinder.flycrispr.neuro.brown.edu). The top and bottom oligos for the gRNA were phosphorylated and annealed using T4 polynucleotide kinase (PNK) (NEB) and then cloned into the pU6 vector (Addgene plasmid #53062), which was linearized with BpiI (NEB). The gRNA and donor DNA were sent to the BestGene for injection into embryos expressing Cas9 in the germline (*y*^*1*^ *M{vas-Cas9.RFP-}ZH-2A w*^*1118*^; RRID: BDSC_55821). To remove the DSRed cassette, transformed flies were balanced and then crossed with flies carrying piggyBac transposase (RRID: BDSC_32073). Flies lacking expression of RFP were recovered and homozygous stocks were sequenced for the *kdm5* locus. The correct removal of the DSRed cassette and presence of the *3xHA* tag was confirmed by PCR sequencing and western blot.

### Immunohistochemistry

For fixation of late 3^rd^ instar larval brain tissue, the CNS was first removed via dissection in cold phosphate buffered saline (PBS, pH 7.4) and allowed to incubate in fixation buffer (4% paraformaldehyde in PBS) at room temperature (RT) for 40-min. After washing for three cycles of 15-min in PBS with 0.2% Triton X-100 (0.2% PBT), brains were incubated with blocking buffer (5% normal goat serum in 0.2% PBT) for 30-min at RT. The brains were then incubated with primary antibodies for two days at 4 °C and washed for three cycles of 15-min in 0.2% PBT. Secondary antibodies were then added and brains were incubated for 2 days at 4 °C. After being washed in PBS for three cycles of 15-min at RT, brains were incubated in a drop of Vectashield mounting medium (Vector Laboratories, H-1000) or DAPI Fluoromount G (SouthernBiotech, OB010020) overnight at 4 °C. Brains were then mounted on glass slides (Superfrost Plus, Fisherbrand), flanked by glass spacers and covered with a final glass coverslip before being used for image analysis.

A similar protocol was followed for immunostaining of pharate adult and 3- to 5-day old adult brain tissue with the following exceptions. Pharate adult heads or whole adult animals were allowed to incubate in fixation buffer (4% paraformaldehyde in 0.2% PBT) at 4 °C for 3-hrs. Heads or whole animals were then washed in 0.2% PBT for three cycles of 15-min at RT. Brains were then dissected from the fixed tissue and blocked for 30 minutes at RT. Antibody incubation and mounting was identical to that described above.

The following primary antibodies were used: mouse anti-Fas2 (1:25, DSHB Cat# 1D4 anti-Fasciclin II, RRID: AB_528235), mouse anti-Trio (1:25, DSHB Cat# 9.4A anti-Trio, RRID: AB_528494), mouse anti-HA (1:50, Cell Signaling Technology Cat# 2367, RRID: AB_10691311), rabbit anti-Prospero (1:1000, Vaessin H; Cell. 1991 Cat# pros, RRID: AB_2569943). Primary antibodies were prepared in 5% NDS/0.2% PBT with 0.02% NaN_3_. The following secondary antibodies were used: goat anti-rabbit (Alexa-488, Alexa 568) and goat anti-mouse (Alexa 488, Alexa 568) (Jackson ImmunoResearch). All secondary antibodies were diluted 1:500 in 5% NDS/0.2% PBT.

### Western Blotting

Western analysis was carried out as previously described (Drelon et al., 2019). Briefly, 3- to 5-day old adult fly heads were homogenized in 2x NuPAGE LDS sample buffer, sonicated for 10 mins, treated with DTT, run on a 4-12% Bis-Tris 1 mm gel and transferred to a PVDF membrane. The following primary antibodies were used: rabbit anti-KDM5 (1:1000, Secombe J; Genes Dev. 2007; Cat# lid, RRID: AB_2569502) and mouse anti-HA (1:1000, Cell Signaling Technology Cat# 2367, RRID: AB_10691311). The following secondary antibodies were used: IRDye^®^ 680RD Donkey anti-Mouse IgG (1:8000; LI-COR Biosciences Cat# 925-68072, RRID: AB_2814912) and IRDye^®^ 800CW Donkey anti-Rabbit IgG (1:8000; LI-COR Biosciences Cat# 926-32213, RRID: AB_621848). Blots were scanned and processed using a LI-COR Odyssey Infrared scanner.

### Image Acquisition

All tissue images were taken on a Leica SP8 confocal microscope using either a 20X air lens (N.A. = 0.75 Air, W.D. = 0.64 mm) or a 63X immersion lens (N.A. = 1.4 Oil, W.D. = 0.14 mm). All MB confocal stacks were taken under either 2X or 2.5X zoom, in a 1024 x 1024 configuration and using 1 μm resolution. Image stacks were processed with Figi (ImageJ). Figures were composed using Microsoft Powerpoint.

### Targeted DamID and Analyses

To profile Pol II occupancy in GMCs and immature neurons of *kdm5*^*140*^ pupae, *kdm5^140^/CyO:gfp; UAS-LT3-dam* or *kdm5^140^/CyO:gfp; UAS-LT3-dam-pol II* flies were crossed with *kdm5^140^/CyO:gfp; R71C09-Gal4* flies and were allowed to lay eggs overnight for 12-14 hrs at 25 °C. Embryos were then moved to 18 °C for a period of 16 days, and GFP negative wandering 3^rd^ instar larvae were transferred to a restrictive temperature of 29 °C for 3 days to induce the expression of the *UAS-dam* and *UAS-dam-pol II* transgenes. As a parallel control, *kdm5* wild-type animals carrying *UAS-LT3-dam* or *UAS-LT3-dam-pol II* were crossed with *R71C09-Gal4* and were also allowed to lay eggs for 24-hrs at 25 °C. Embryos were then moved to 18 °C for 9 days and subsequently transferred to 29 °C for 3 days.

For *kdm5* knockdown experiments, flies carrying either *UAS-LT3-dam; UAS-kdm5_shRNA* or *UAS-LT3-dam-pol II; UAS-kdm5_shRNA* were crossed with flies carrying *UAS-tubulin-Gal80^ts^; R71C09-Gal4*. Parallel crosses were also performed with flies bearing a scrambled shRNA sequence in place of *kdm5* shRNA. All flies were allowed to lay eggs overnight for 12-14 hrs at 25 °C. Embryos were then transferred to 18 °C for 4 days and then subsequently transferred to 29 °C for 6 days.

Tissue processing for all TaDa experiments was performed as previously described in Marshall et al., 2016 with the following modifications. A total of 40 pupae of each genotype were homogenized in 500 mM EDTA followed by DNA extraction using the Zymo Quick-DNA Miniprep Plus Kit. DpnI digestion, PCR adaptor ligation, DpnII digestion, and PCR amplification were performed as described. DNA was sonicated using a Diagenode Bioruptor for 8-10 cycles (5-min at high power, 30-s on/30-s off) and analyzed using an Agilent Bioanalyzer. DamID adaptor removal and DNA cleanup were performed as previously described (Marshall et al., 2016) and samples were submitted to BGI for sequencing.

Sequencing libraries were prepared at BGI Genomics following a ChIP-seq workflow. DNA fragments were first end-repaired and dA-tailed using End Repair and A-Tailing (ERAT) enzyme. Adaptors were then ligated for sequencing and ligated DNA purified using AMPure beads. DNA was then PCR amplified with BGI primers for 8 cycles and PCR purified with AMPure beads. DNA was then homogenized, circularized, digested, and again purified. DNA was then prepared into proprietary DNA nanoballs (DNB™) for sequencing on a BGISEQ-500 platform with 50 bp read length and 20 M clean reads.

### *Drosophila* dSO Avoidance Assays

*Drosophila* dSO avoidance assays were carried out as previously described with modifications (Fernandez et al., 2014; Suh et al., 2004). Briefly, 24 hours before the experiment 100 wild-type adult flies (“emitters”) as well as 60-65 experimental or control flies (“responders”) were each transferred to standard food vials and housed at 25 °C. For each experimental or control cohort, 100 emitters were transferred to an empty tube (“dSO vial”) and vortexed at maximum speed in 15-s bouts, at 5-s intervals, for 1-min and then removed. The dSO vial was immediately cleared and placed in one arm of a T-maze (CelExplorer Labs) and an empty vial in another. Responders were then deposited into the T-maze elevator and allowed to rest for 1-min. Responders were then given 1-min to choose between the dSO vial and the empty vial. The locations of the dSO and empty vials were alternated between trials to control for potential environmental confounders. Additionally, a separate cohort of emitters was used for each trial. The Performance Index (“PI”) was calculated by subtracting the number of flies within the dSO vial from those within the empty vial and dividing by the total number of flies.

### QUANTIFICATION AND STATISTICAL ANALYSES

#### Statistical Analyses

For MB morphological analyses, results are presented as bar plots for which percentage of brains with MB lobe growth and/or guidance defects are calculated. For these analysis, N = number of brains examined. “Growth defects” were defined by an overgrown, stunted, or absent lobe and “guidance defects” were defined by a full or partially misprojected lobe. In the case where both defect types were observed in a single brain, the defect was categorized as a “growth and guidance defect.” All MB statistical analysis was performed using GraphPad Prism 8.4 (GraphPad Software, Inc., Ca, US). A 2 x N contingency table was used when comparing MB defects of more than two genotypes, where N = number of genotypes, and significance was determined using a chi-square test with either a Yates’ or Bonferroni correction with * = *P* < 0.05, ** = *P* < 0.01, *** = *P* < 0.001, and **** = *P* < 0.0001.

For *Drosophila* dSO Avoidance Assays, results are presented as Performance Index (PI) with mean ± SEM. Significance was determined using a one-way ANOVA with Tukey post-hoc with * = *P* < 0.05 and ** = *P* < 0.01. Statistical analysis was performed using GraphPad Prism.

Gene Ontology enrichment analysis was done using PANTHER Overrepresentation testing (http://geneontology.org/; Mi et al., 2013) with a Fisher’s exact test (FDR < 0.05). Annotation version and release date: GO Ontology database DOI: 10.5281/zenodo.3873405, Released 2020-06-01.

For Targeted DamID analyses, sequencing data were aligned to release 6 of the *Drosophila melanogaster* genome and processed using damidseq_pipeline as previously described (Marshall and Brand, 2017; Marshall et al., 2016). RNA Pol II occupancy over gene bodies was calculated via polii.gene.call (Marshall et al., 2016). Differentially expressed genes were called via the NOIseq R package (Tarazona et al., 2011); briefly, RNA pol II gene occupancy scores were scaled, and inverse log values used as input to NOIseq with parameters of upper quantile normalization and biological replicates. DEGs were called with a q value of 0.95.

For CATaDa analyses, Dam-only BAM files generated via damidseq_pipeline were converted to 75 nt bins via bam2coverage, and replicates averaged. Peaks were called separately on the wild-type and mutant conditions via find_peaks (Marshall et al., 2016) with a minimum quantile of 0.95, before combining and merging peaks with BEDTools (Quinlan and Hall, 2010), and plotting heatmaps with SeqPlots (Stempor and Ahringer, 2016). Average Dam occupancy values over peaks for each biological replicate was determined via polii.gene.call and differential occupancy called via NOIseq as with RNA Pol II occupancy above. Peaks called as significant were associated with the nearest gene promoter via peaks2genes (Marshall et al., 2016).

## SUPPLEMENTAL FIGURE TITLES AND LEGENDS

**Figure S1.**
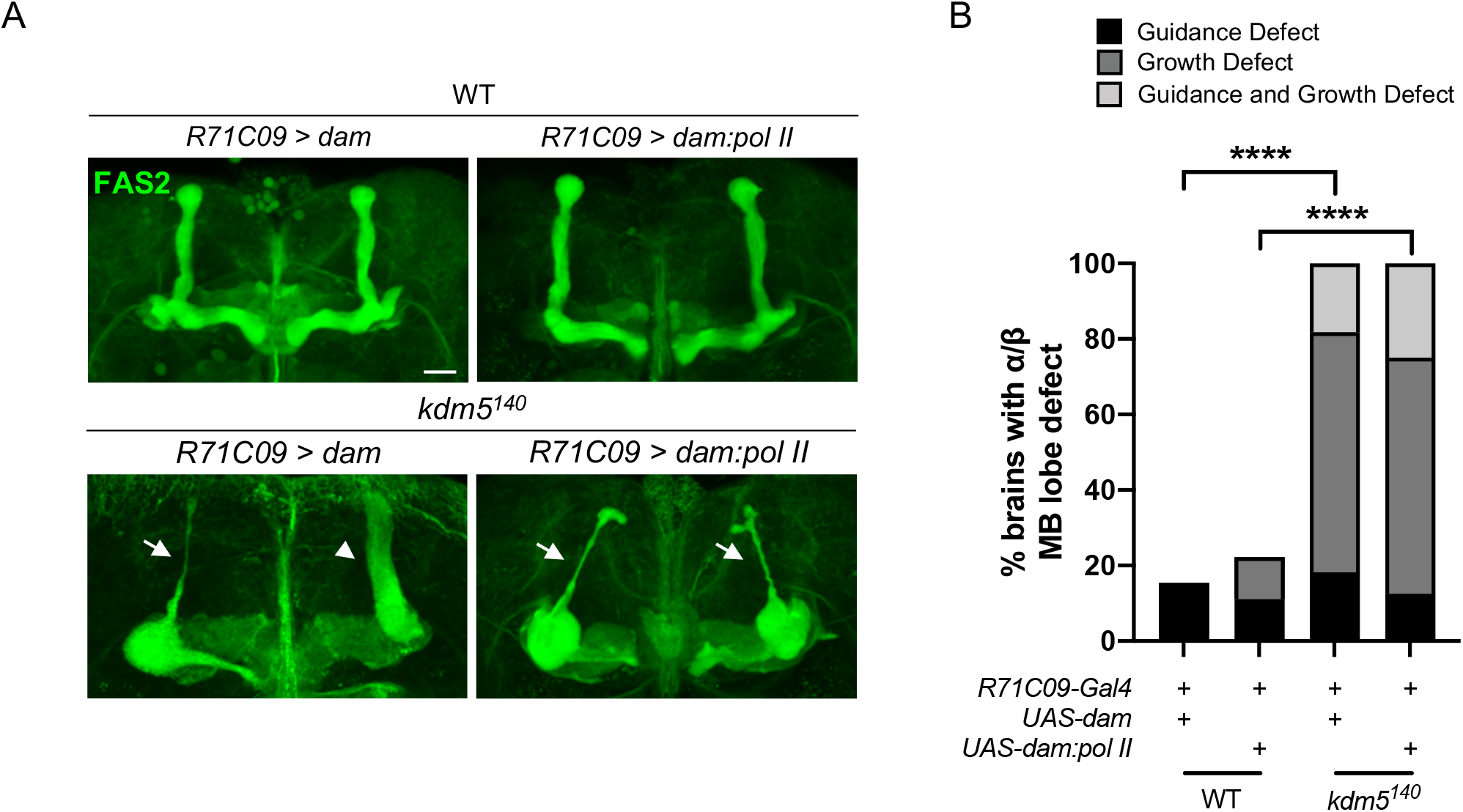
*kdm5*^*140*^ Pharate Adults Expressing TaDa Genetic Elements Present with Significant MB Morphological Defects. Related to Figure 5. (A) Representative α/β lobe Z projections of pharate *kdm5*^*140*^ animals expressing TaDa genetic elements and their respective controls. The antibody anti-Fas2 is used to visualize α/β lobes. Arrows indicate growth defects. Scale bar represents 20 μm. (B) Quantification of α/β lobe defects in pharate *kdm5*^*140*^ animals expressing TaDa genetic elements and their respective controls. *n* = 8-22 (mean *n* = 13). **** *P* < 0.0001 (chi-square test with Bonferroni Correction).

**Figure S2.**
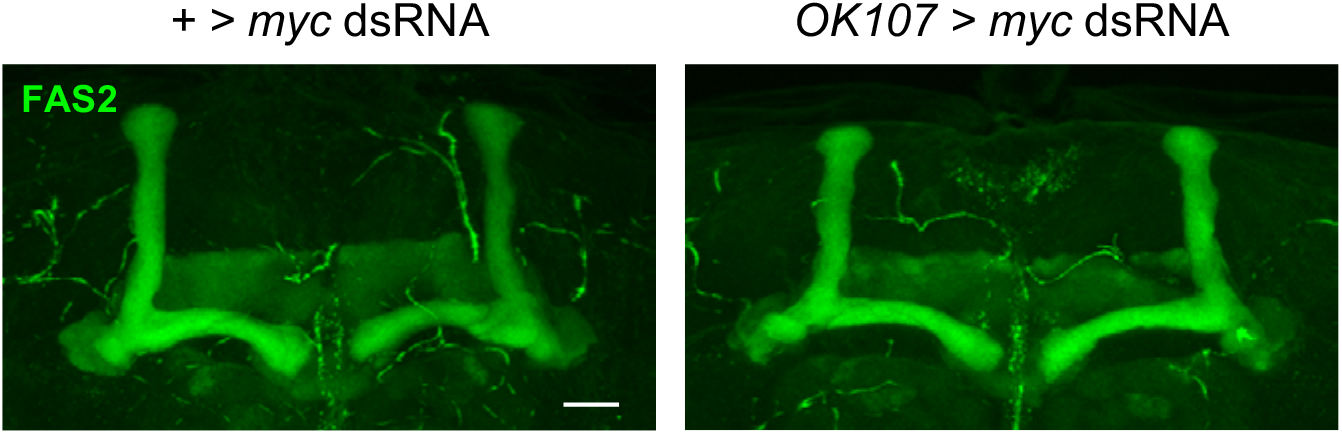
Knockdown of *myc* within Kenyon Cells Does Not Result in Morphological Defects of the MB. Related to Figure 5. Representative α/β lobe Z projections of adults expressing *myc* dsRNA driven by OK107-Gal4. No gross morphological defects were detected in the KD or control conditions *n* = 29-30.

**Figure S3.**
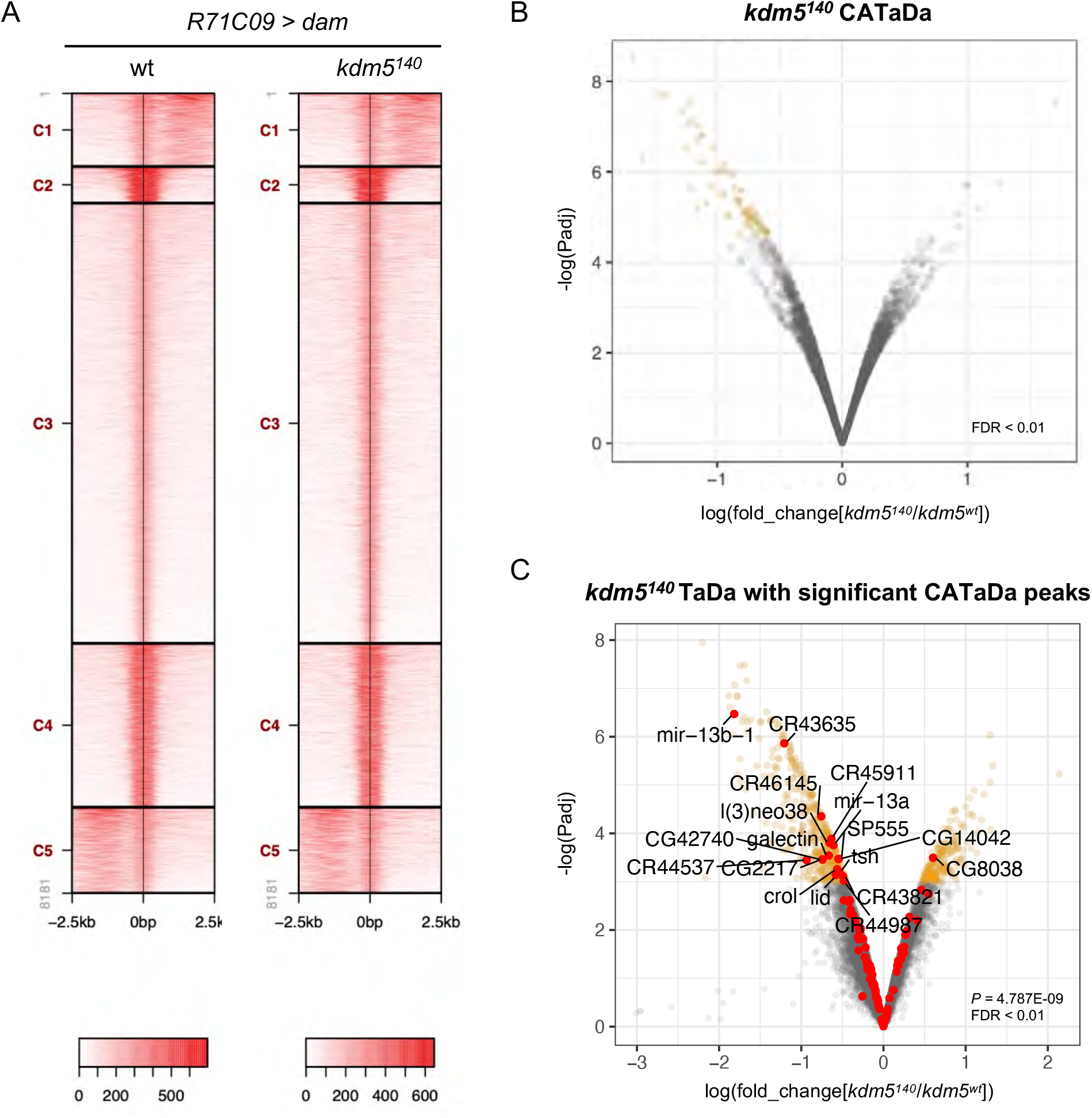
CATaDa Analyses Reveal Minimal Changes to Chromatin Accessibility within GMCs and Immature Neurons Upon Loss of *kdm5*. Related to Figure 5. (A) Representative CATaDa profiles of *R71C09-Gal4* expressing cells for *kdm5*^*140*^ and wildtype. Heat maps show Dam binding profiles for the greatest 8181 peaks for each genotype and are organized into five clusters. (B) Volcano plot showing changes to chromatin accessibility within GMCs and immature neurons of *kdm5*^*140*^ animals compared to wildtype. Genes with an FDR < 0.01 are in yellow. CATaDa analyses were performed in quintuplicate. (C) Volcano plot showing changes to chromatin accessibility for DEGs from *kdm5*^*140*^ TaDa. In yellow are *kdm5*^*140*^ TaDa DEGs with an FDR < 0.05. In red are *kdm5*^*140*^ TaDa DEGs with associated changes in chromatin accessibility for FDR < 0.01 (Fisher’s exact test, *P* = 4.79E-09).

**Figure S4.**
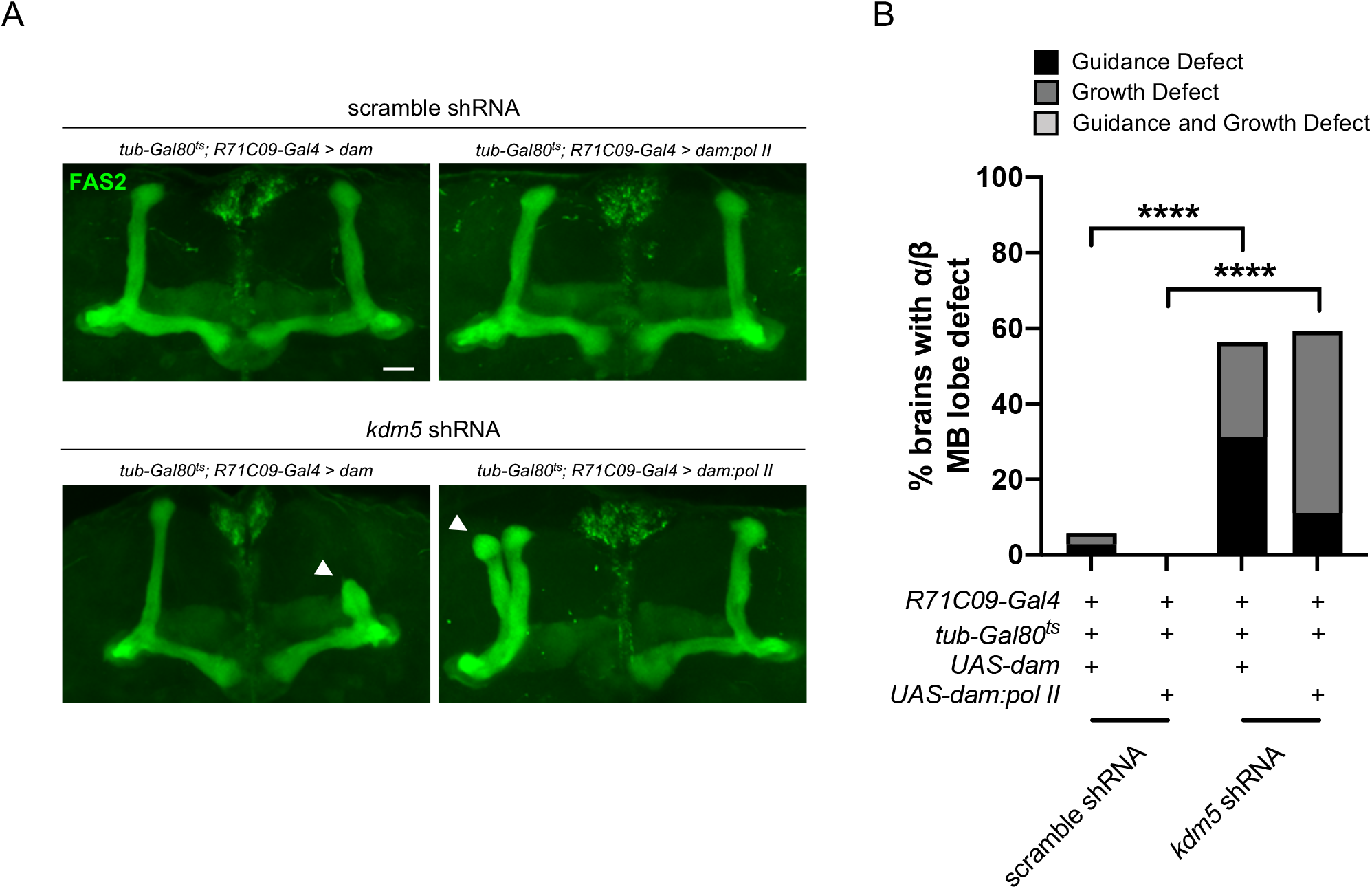
Adults Expressing *R71C09-Gal4* driven *kdm5* shRNA in Tandem with TaDa Genetic Elements Present with Significant MB Morphological Defects. Related to Figure 5. Representative α/β lobe Z projections of adult *R71C09 > kdm5* shRNA animals expressing TaDa genetic elements and their respective controls. The antibody anti-Fas2 is used to visualize α/β lobes. Arrowheads indicate guidance defects. Scale bar represents 20 μm. (B) Quantification of α/β lobe Z projections of adult *R71C09 > kdm5* shRNA animals expressing TaDa genetic elements and their respective controls. *n* = 16-34 (mean *n* = 28). **** *P* < 0.0001 (chi-square test with Bonferroni Correction).

**Figure S5.**
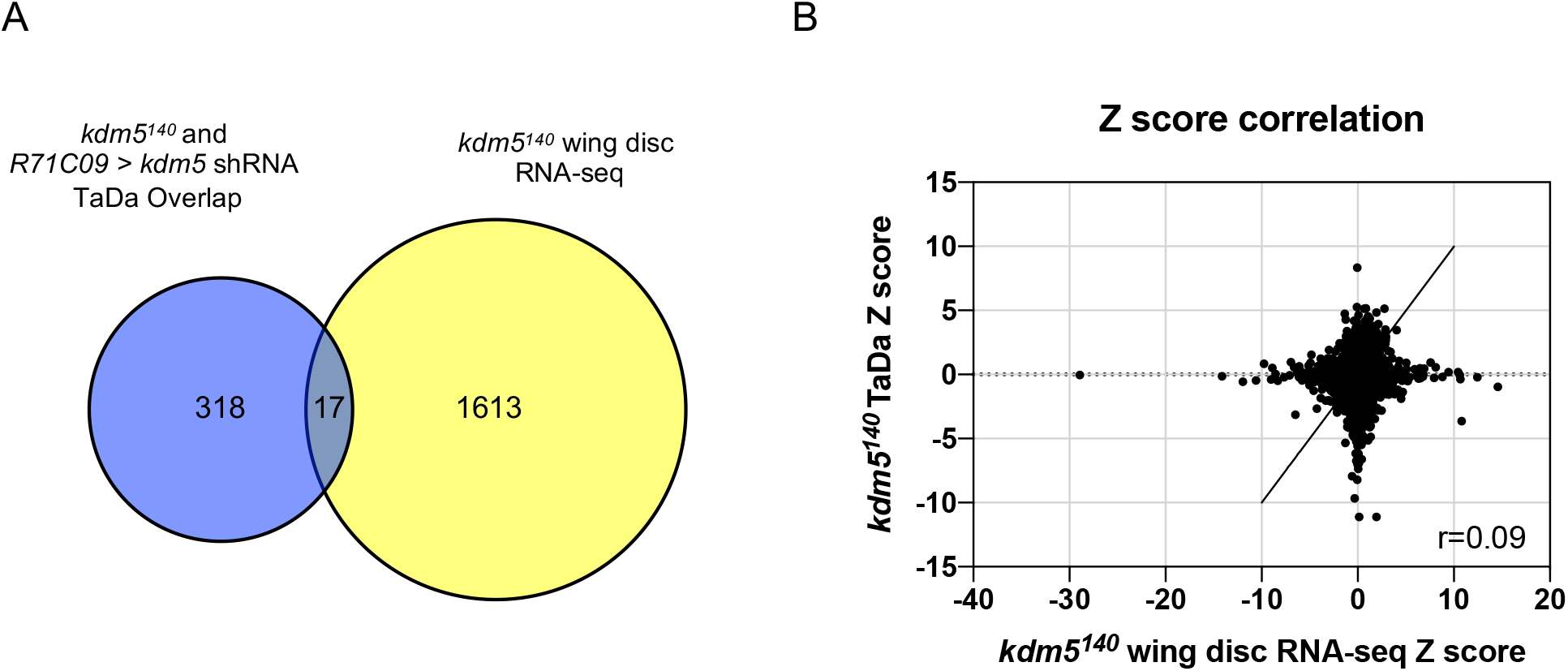
Transcriptome Analyses Reveal that KDM5-Regulated Gene Expression is Tissue-Specific. Related to Figure 5. (A) Venn diagram illustrating intersection of *kdm5*^*140*^ and *kdm5* shRNA overlapping TaDa data with that from a previously published *kdm5*^*140*^ mRNA-seq wing disc dataset (Drelon et al., 2018). (B) Correlation of Z scores between *kdm5*^*140*^ TaDa and *kdm5*^*140*^ wing disc mRNA-seq (Drelon et al., 2018) datasets. Pearson’s R = 0.09.

**Table S1. DEGs from *kdm5*^*140*^ TaDa. Related to Figure 5.**

**Table S2. GO Categories from *kdm5*^*140*^ TaDa. Related to Figure 5**

**Table S3. DEGs from *kdm5* shRNA TaDa. Related to Figure 5.**

**Table S4. GO Categories from *kdm5* shRNA TaDa. Related to Figure 5.**

**Table S5. Overlapping DEGs between TaDa and *kdm5*^*140*^ Wing Disc RNA-seq Datasets. Related to Figure 5.**

**Table S6. TaDa DEGs that are Direct Dam:Pros Targets. Related to Figure 7.**

## REFERENCES

1. Androschuk, A., Al-Jabri, B., and Bolduc, F. V. (2015). From learning to memory: What flies can tell us about intellectual disability treatment. Front. Psychiatry 6, 1–16.

2. Androschuk, A., He, R.X., Weber, S., Rosenfelt, C., and Bolduc, F.V. (2018). Stress Odorant Sensory Response Dysfunction in Drosophila Fragile X Syndrome Mutants. Front. Mol. Neurosci. 11, 1–11.

3. Aso, Y., Hattori, D., Yu, Y., Johnston, R.M., Iyer, N.A., Ngo, T.T.B., Dionne, H., Abbott, L.F., Axel, R., Tanimoto, H., et al. (2014). The neuronal architecture of the mushroom body provides a logic for associative learning. Elife 3, e04577.

4. Aughey, G.N., Estacio Gomez, A., Thomson, J., Yin, H., and Southall, T.D.(2018). CATaDa reveals global remodelling of chromatin accessibility during stem cell differentiation in vivo. Elife 7, 1–22.

5. Aughey, G.N., Cheetham, S.W., and Southall, T.D.(2019). DamID as a versatile tool for understanding gene regulation. Development 146, dev173666.

6. Bates, K.E., Sung, C.S., and Robinow, S. (2010). The *unfulfilled* gene is required for the development of mushroom body neuropil in *Drosophila*. Neural Dev. 5, 4.

7. Bellosta, P., and Gallant, P. (2010). Myc function in Drosophila. Genes and Cancer 1, 542–546.

8. Bhaumik, S.R., Smith, E., and Shilatifard, A. (2007). Covalent modifications of histones during development and disease pathogenesis. Nat. Struct. Mol. Biol. 14, 1008–1016.

9. van Bokhoven, H. (2011). Genetic and Epigenetic Networks in Intellectual Disabilities. Annu. Rev. Genet. 45, 81–104.

10. Brookes, E., Laurent, B., Õunap, K., Carroll, R., John, B., Field, M., Schwartz, C.E., Gecz, J., and Shi, Y. (2015). Mutations in the intellectual disability gene KDM5C reduce protein stability and demethylase activity. 24, 2861–2872.

11. Chen, K., Luan, X., Liu, Q., Wang, J., Chang, X., Snijders, A.M., Mao, J.-H., Secombe, J., Dan, Z., Chen, J.-H., et al. (2019). Drosophila Histone Demethylase KDM5 Regulates Social Behavior through Immune Control and Gut Microbiota Maintenance. Cell Host Microbe 1–16.

12. Choksi, S.P., Southall, T.D., Bossing, T., Edoff, K., de Wit, E., Fischer, B.E.E., van Steensel, B., Micklem, G., and Brand, A.H.(2006). Prospero Acts as a Binary Switch between Self-Renewal and Differentiation in Drosophila Neural Stem Cells. Dev. Cell 11, 775–789.

13. Doe, C.Q., Chu-LaGraff, Q., Wright, D.M., and Scott, M.P.(1991). The prospero gene specifies cell fates in the drosophila central nervous system. Cell 65, 451–464.

14. Drelon, C., Belalcazar, H.M., and Secombe, J. (2018). The histone demethylase KDM5 is essential for larval growth in Drosophila. 209, 773–787.

15. Drelon, C., Rogers, M.F., Belalcazar, H.M., and Secombe, J. (2019). The histone demethylase KDM5 controls developmental timing in Drosophila by promoting prothoracic gland endocycles. Dev. 146.

16. Dubnau, J., Grady, L., Kitamoto, T., and Tully, T. (2001). Disruption of neurotransmission in Drosophila mushroom body blocks retrieval but not acquisition of memory. Nature 411, 476–480.

17. Elbert, A., and Bérubé, N.G. (2014). Chromatin Structure and Intellectual Disability Syndromes. Neurosci. Biobehav. Rev 46, 161–174.

18. Fenckova, M., Blok, L.E.R., Asztalos, L., Goodman, D.P., Cizek, P., Singgih, E.L., Glennon, J.C., IntHout, J., Zweier, C., Eichler, E.E., et al. (2019). Habituation Learning Is a Widely Affected Mechanism in Drosophila Models of Intellectual Disability and Autism Spectrum Disorders. Biol. Psychiatry.

19. Fernandez, R.W.et al. (2017). Modulation of social space by dopamine in *Drosophila melanogaster*, but no effect on the avoidance of the *Drosophila* stress odorant Supplementary material. Biol. Lett.

20. Fernandez, R.W., Nurilov, M., Feliciano, O., McDonald, I.S., and Simon, A.F.(2014). Straightforward assay for quantification of social avoidance in Drosophila melanogaster. J. Vis. Exp. 1–8.

21. Froldi, F., Szuperak, M., Weng, C.F., Shi, W., Papenfuss, A.T., and Cheng, L.Y.(2015). The transcription factor Nerfin-1 prevents reversion of neurons into neural stem cells. Genes Dev. 29, 129–143.

22. Furukubo-Ttokunaga, K., Aadachi, Y., Kurusu, M., and Walldorf, U. (2009). Brain patterning defects caused by mutations of the twin of eyeless gene in Drosophila melanogaster. Fly (Austin). 3.

23. Gajan, A., Barnes, V.L., Liu, M., Saha, N., and Pile, L.A.(2016). The histone demethylase dKDM5/LID interacts with the SIN3 histone deacetylase complex and shares functional similarities with SIN3. Epigenetics and Chromatin 9, 1–23.

24. Gombos, R., Migh, E., Antal, O., Mukherjee, A., Jenny, A., and Mihaly, J. (2015). The Formin DAAM Functions as Molecular Effector of the Planar Cell Polarity Pathway during Axonal Development in Drosophila. J. Neurosci. 35, 10154–10167.

25. Gonçalves, T.F., Gonçalves, A.P., Fintelman Rodrigues, N., dos Santos, J.M., Pimentel, M.M.G., and Santos-Rebouças, C.B.(2014). KDM5C mutational screening among males with intellectual disability suggestive of X-Linked inheritance and review of the literature. Eur. J. Med. Genet. 57, 138–144.

26. Greer, E.L., and Shi, Y. (2012). Histone methylation: A dynamic mark in health, disease and inheritance. Nat. Rev. Genet. 13, 343–357.

27. Grewal, S.S., Li, L., Orian, A., Eisenman, R.N., and Edgar, B.A.(2005). Myc-dependent regulation of ribosomal RNA synthesis during Drosophila development. Nat. Cell Biol. 7, 295–302.

28. Harmeyer, K.M., Facompre, N.D., Herlyn, M., and Basu, D. (2017). JARID1 Histone Demethylases: Emerging Targets in Cancer. Trends in Cancer 3, 713–725.

29. Hattori, D., Demir, E., Kim, H.W., Viragh, E., Zipursky, S.L., and Dickson, B.J.(2007). Dscam diversity is essential for neuronal wiring and self-recognition. Nature 449, 223–227.

30. Heisenberg, M., and Technau, G. (1982). Neural reorganization during metamorphosis of the corpora pedunculata in Drosophila melanogaster. Nature 295, 405–407.

31. Heisenberg, M., Borst, A., Wagner, S., and Byers, D. (1985). Drosophila mushroom body mutants are deficient in olfactory learning. J. Neurogenet. 2, 1–30.

32. Ito, K., and Hotta, Y. (1992). Proliferation pattern of postembryonic neuroblasts in the brain of Drosophila melanogaster. Dev. Biol. 149, 134–148.

33. Iwase, S., Brookes, E., Agarwal, S., Badeaux, A.I., Ito, H., Vallianatos, C.N., Tomassy, G.S., Kasza, T., Lin, G., Thompson, A., et al. (2016a). A Mouse Model of X-linked Intellectual Disability Associated with Impaired Removal of Histone Methylation. Cell Rep. 14, 1000–1009.

34. Iwase, S., Brookes, E., Agarwal, S., Egan, B., Xu, J., Iwase, S., Brookes, E., Agarwal, S., Badeaux, A.I., Ito, H., et al. (2016b). A Mouse Model of X-linked Intellectual Disability Associated with Impaired Removal of Histone Report A Mouse Model of X-linked Intellectual Disability Associated with Impaired Removal of Histone Methylation. CellReports 14, 1000–1009.

35. Kelly, S.M., Bienkowski, R., Banerjee, A., Melicharek, D.J., Brewer, Z.A., Marenda, D.R., Corbett, A.H., and Moberg, K.H.(2016). The Drosophila ortholog of the Zc3h14 RNA binding protein acts within neurons to pattern axon projection in the developing brain. Dev. Neurobiol. 76, 93–106.

36. Kochinke, K., Zweier, C., Nijhof, B., Fenckova, M., Cizek, P., Honti, F., Keerthikumar, S., Oortveld, M.A.W., Kleefstra, T., Kramer, J.M., et al. (2016). Systematic Phenomics Analysis Deconvolutes Genes Mutated in Intellectual Disability into Biologically Coherent Modules. Am. J. Hum. Genet. 98, 149–164.

37. Kong, S.-Y., Kim, W., Lee, H.-R., and Kim, H.-J. (2017). The histone demethylase KDM5A is required for the repression of astrocytogenesis and regulated by the translational machinery in neural progenitor cells. FASEB J. fj.201700780R.

38. Kuzin, A., Brody, T., Moore, A.W., and Odenwald, W.F.(2005). Nerfin-1 is required for early axon guidance decisions in the developing Drosophila CNS. Dev. Biol. 277, 347–365.

39. Lee, M.G., Norman, J., Shilatifard, A., and Shiekhattar, R. (2007). Physical and Functional Association of a Trimethyl H3K4 Demethylase and Ring6a/MBLR, a Polycomb-like Protein. Cell 128, 877–887.

40. Lee, N., Erdjument-Bromage, H., Tempst, P., Jones, R.S., and Zhang, Y. (2009). The H3K4 Demethylase Lid Associates with and Inhibits Histone Deacetylase Rpd3. Mol. Cell. Biol. 29, 1401–1410.

41. Leonard, H., and Wen, X. (2002). The epidemiology of mental retardation: Challenges and opportunities in the new millennium. Ment. Retard. Dev. Disabil. Res. Rev. 8, 117–134.

42. Li, G., Ruan, X., Auerbach, R.K., Sandhu, K.S., Zheng, M., Wang, P., Poh, H.M., Goh, Y., Lim, J., Zhang, J., et al. (2012). Extensive promoter-centered chromatin interactions provide a topological basis for transcription regulation. Cell 148, 84–98.

43. Li, H.H., Kroll, J.R., Lennox, S.M., Ogundeyi, O., Jeter, J., Depasquale, G., and Truman, J.W.(2014). A GAL4 driver resource for developmental and behavioral studies on the larval CNS of Drosophila. Cell Rep. 8, 897–908.

44. Liefke, R., Oswald, F., Alvarado, C., Ferres-Marco, D., Mittler, G., Rodriguez, P., Dominguez, M., and Borggrefe, T. (2010). Histone demethylase KDM5A is an integral part of the core Notch-RBP-J repressor complex. Genes Dev. 24, 590–601.

45. Liu, X., and Secombe, J. (2015). The Histone Demethylase KDM5 Activates Gene Expression by Recognizing Chromatin Context through Its PHD Reader Motif. Cell Rep. 13, 2219–2231.

46. Liu, X., Greer, C., and Secombe, J. (2014a). KDM5 Interacts with Foxo to Modulate Cellular Levels of Oxidative Stress. PLoS Genet. 10.

47. Liu, X., Greer, C., and Secombe, J. (2014b). KDM5 Interacts with Foxo to Modulate Cellular Levels of Oxidative Stress. PLoS Genet. 10.

48. Liu, X., Shen, J., Xie, L., Wei, Z., Wong, C., Li, Y., Zheng, X., and Li, P. (2020). Mitotic Implantation of the Transcription Factor Prospero via Phase Separation Drives Terminal Neuronal Differentiation Article Mitotic Implantation of the Transcription Factor Prospero via Phase Separation Drives Terminal Neuronal Differentiation. Dev. Cell 1–17.

49. Lloret-Llinares, M., Pérez-Lluch, S., Rossell, D., Morán, T., Ponsa-Cobas, J., Auer, H., Corominas, M., and Azorín, F. (2012). DKDM5/LID regulates H3K4me3 dynamics at the transcription-start site (TSS) of actively transcribed developmental genes. Nucleic Acids Res. 40, 9493–9505.

50. Lopez-Bigas, N., Kisiel, T.A., DeWaal, D.C., Holmes, K.B., Volkert, T.L., Gupta, S., Love, J., Murray, H.L., Young, R.A., and Benevolenskaya, E. V. (2008). Genome-wide Analysis of the H3K4 Histone Demethylase RBP2 Reveals a Transcriptional Program Controlling Differentiation. Mol. Cell 31, 520–530.

51. Lussi, Y.C., Mariani, L., Friis, C., Peltonen, J., Myers, T.R., Krag, C., Wong, G., and Salcini, A.E.(2016). Impaired removal of H3K4 methylation affects cell fate determination and gene transcription. Dev. 143, 3751–3762.

52. Marchetti, G., and Tavosanis, G. (2017). Steroid Hormone Ecdysone Signaling Specifies Mushroom Body Neuron Sequential Fate via Chinmo. Curr. Biol. 27, 3017–3024.e4.

53. Mariani, L., Lussi, Y.C., Vandamme, J., Riveiro, A., and Salcini, A.E.(2016). The H3K4me3/2 histone demethylase RBR-2 controls axon guidance by repressing the actin-remodeling gene wsp-1. Development 143, 851–863.

54. Marshall, O.J., and Brand, A.H.(2017). Chromatin state changes during neural development revealed by in vivo cell-type specific profiling. Nat. Commun. 8, 1–9.

55. Marshall, O.J., Southall, T.D., Cheetham, S.W., and Brand, A.H.(2016). Cell-type-specific profiling of protein-DNA interactions without cell isolation using targeted DamID with next-generation sequencing. Nat. Protoc. 11, 1586–1598.

56. Mi, H., Muruganujan, A., Casagrande, J.T., and Thomas, P.D.(2013). Large-scale gene function analysis with the panther classification system. Nat. Protoc. 8, 1551–1566.

57. Michel, C.I.(2004). Defective Neuronal Development in the Mushroom Bodies of Drosophila Fragile X Mental Retardation 1 Mutants. J. Neurosci. 24, 5798–5809.

58. Morris, J.H., Apeltsin, L., Newman, A.M., Baumbach, J., Wittkop, T., Su, G., Bader, G.D., and Ferrin, T.E.(2011). ClusterMaker: A multi-algorithm clustering plugin for Cytoscape. BMC Bioinformatics 12, 1–14.

59. Najmabadi, H., Hu, H., Garshasbi, M., Zemojtel, T., Abedini, S.S., Chen, W., Hosseini, M., Behjati, F., Haas, S., Jamali, P., et al. (2011). Deep sequencing reveals 50 novel genes for recessive cognitive disorders. Nature 478, 57–63.

60. Navarro-Costa, P., McCarthy, A., Prudêncio, P., Greer, C., Guilgur, L.G., Becker, J.D., Secombe, J., Rangan, P., and Martinho, R.G.(2016). Early programming of the oocyte epigenome temporally controls late prophase i transcription and chromatin remodelling. Nat. Commun. 7.

61. Nishibuchi, G., Shibata, Y., Hayakawa, T., Hayakawa, N., Ohtani, Y., Sinmyozu, K., Tagami, H., and Nakayama, J.I.(2014). Physical and functional interactions between the histone H3K4 demethylase KDM5A and the nucleosome remodeling and deacetylase (NuRD) complex. J. Biol. Chem. 289, 28956–28970.

62. van Oevelen, C., Wang, J., Asp, P., Yan, Q., Kaelin, W.G., Kluger, Y., and Dynlacht, B.D.(2008). A Role for Mammalian Sin3 in Permanent Gene Silencing. Mol. Cell 32, 359–370.

63. Park, S., Kim, G.W., Kwon, S.H., and Lee, J.S.(2020). Broad domains of histone H3 lysine 4 trimethylation in transcriptional regulation and disease. FEBS J. 287, 2891–2902.

64. Parkel, S., Lopez-Atalaya, J.P., and Barco, A. (2013). Histone H3 lysine methylation in cognition and intellectual disability disorders. Learn. Mem. 20, 570–579.

65. Quinlan, A.R., and Hall, I.M.(2010). BEDTools: A flexible suite of utilities for comparing genomic features. Bioinformatics 26, 841–842.

66. Ropers, H.H.(2010). Genetics of Early Onset Cognitive Impairment. Annu. Rev. Genomics Hum. Genet. 11, 161–187.

67. Ropers, H.-H., and Hamel, B.C.J. (2005). X-linked mental retardation. Nat. Rev. Genet. 6, 46–57.

68. Scandaglia, M., Lopez-Atalaya, J.P., Medrano-Fernandez, A., Lopez-Cascales, M.T., del Blanco, B., Lipinski, M., Benito, E., Olivares, R., Iwase, S., Shi, Y., et al. (2017). Loss of Kdm5c Causes Spurious Transcription and Prevents the Fine-Tuning of Activity-Regulated Enhancers in Neurons. Cell Rep. 21, 47–59.

69. Secombe, J., Li, L., Carlos, L., and Eisenman, R.N.(2007). The Trithorax group protein Lid is a trimethyl histone H3K4 demethylase required for dMyc-induced cell growth. Genes Dev. 21, 537–551.

70. Shannon, P., Markiel, A., Ozier, O., Baliga, N.S., Wang, J.T., Ramage, D., Amin, N., Schwikowski, B., and Ideker, T. (2003). Cytoscape: A Software Environment for Integrated Models of Biomolecular Interaction Networks. Genome Res. 13, 2498–2504.

71. Southall, T.D., Gold, K.S., Egger, B., Davidson, C.M., Caygill, E.E., Marshall, O.J., and Brand, A.H.(2013). Cell-type-specific profiling of gene expression and chromatin binding without cell isolation: Assaying RNA pol II occupancy in neural stem cells. Dev. Cell 26, 101–112.

72. Stempor, P., and Ahringer, J. (2016). SeqPlots - Interactive software for exploratory data analyses, pattern discovery and visualization in genomics. Wellcome Open Res. 1, 14.

73. Suh, G.S.B., Wong, A.M., Hergarden, A.C., Wang, J.W., Simon, A.F., Benzer, S., Axel, R., and Anderson, D.J.(2004). A single population of olfactory sensory neurons mediates an innate avoidance behaviour in Drosophila. Nature 431, 854–859.

74. Syed, M.H., Mark, B., and Doe, C.Q.(2017). Steroid hormone induction of temporal gene expression in drosophila brain neuroblasts generates neuronal and glial diversity. Elife 6, 1–23.

75. Tarazona, S., García-alcalde, F., Dopazo, J., Ferrer, A., and Conesa, A. (2011). Differential expression in RNA-seq: A matter of depth. Genome Res. 21, 2213–2223.

76. Tea, J.S., Chihara, T., and Luo, L. (2010). Histone deacetylase Rpd3 regulates olfactory projection neuron dendrite targeting via the transcription factor prospero. J. Neurosci. 30, 9939–9946.

77. The Gene Ontology Consortium (2000). Gene Ontology: tool for the unification of biology. Gene Expr. 25, 25–29.

78. The Gene Ontology Consortium (2019). The Gene Ontology Resource: 20 years and still GOing strong. Nucleic Acids Res. 47, D330–D338.

79. Thurman, R.E., Rynes, E., Humbert, R., Vierstra, J., Maurano, M.T., Haugen, E., Sheffield, N.C., Stergachis, A.B., Wang, H., Vernot, B., et al. (2012). The accessible chromatin landscape of the human genome. Nature 489, 75–82.

80. Truman, J.W., and Bate, M. (1988). Spatial and temporal patterns of neurogenesis in the central nervous system of Drosophila melanogaster. Dev. Biol. 125, 145–157.

81. Vaessin, H., Grell, E., Wolff, E., Bier, E., Jan, L.Y., and Jan, Y.N.(1991). prospero is expressed in neuronal precursors and encodes a nuclear protein that is involved in the control of axonal outgrowth in Drosophila. Cell 67, 941–953.

82. Vallianatos, C.N., and Iwase, S. (2015). Disrupted intricacy of histone H3K4 methylation in neurodevelopmental disorders. Epigenomics 7, 503–519.

83. Vallianatos, C.N., Farrehi, C., Friez, M.J., Burmeister, M., Keegan, C.E., and Iwase, S. (2018). Altered Gene-Regulatory Function of KDM5C by a Novel Mutation Associated With Autism and Intellectual Disability. Front. Mol. Neurosci. 11, 1–12.

84. Vallianatos, C.N., Raines, B., Porter, R.S., Bonefas, K.M., Wu, M.C., Garay, P.M., Collette, K.M., Seo, Y.A., Dou, Y., Keegan, C.E., et al. (2020). Mutually suppressive roles of KMT2A and KDM5C in behaviour, neuronal structure, and histone H3K4 methylation. Commun. Biol. 3.

85. Wei, G., Deng, X., Agarwal, S., Iwase, S., Disteche, C., and Xu, J. (2016). Patient Mutations of the Intellectual Disability Gene KDM5C Downregulate Netrin G2 and Suppress Neurite Growth in Neuro2a Cells. J. Mol. Neurosci.

86. Wise, A., Tenezaca, L., Fernandez, R.W., Schatoff, E., Flores, J., Ueda, A., Zhong, X., Wu, C.F., Simon, A.F., and Venkatesh, T. (2015). Drosophila mutants of the autism candidate gene neurobeachin (rugose) exhibit neuro-developmental disorders, aberrant synaptic properties, altered locomotion, and impaired adult social behavior and activity patterns. J. Neurogenet. 29, 135–143.

87. Witteveen, J.S., Willemsen, M.H., Dombroski, T.C.D., Van Bakel, N.H.M., Nillesen, W.M., Van Hulten, J.A., Jansen, E.J.R., Verkaik, D., Veenstra-Knol, H.E., Van Ravenswaaij-Arts, C.M.A., et al. (2016). Haploinsufficiency of MeCP2-interacting transcriptional co-repressor SIN3A causes mild intellectual disability by affecting the development of cortical integrity. Nat. Genet. 48, 877–887.

88. Yuan, L., Hu, S., Okray, Z., Ren, X., De Geest, N., Claeys, A., Yan, J., Bellefroid, E., Hassan, B.A., and Quan, X.J.(2016). The Drosophila neurogenin tap functionally interacts with the Wnt-PCP pathway to regulate neuronal extension and guidance. Dev. 143, 2760–2766.

89. Zamurrad, S., Hatch, H.A.M., Drelon, C., Belalcazar, H.M., and Secombe, J. (2018). A Dro`used by Mutations in the Histone Demethylase KDM5. Cell Rep. 22, 2359–2369.

90. Zhan, X.L., Clemens, J.C., Neves, G., Hattori, D., Flanagan, J.J., Hummel, T., Vasconcelos, M.L., Chess, A., and Zipursky, S.L.(2004). Analysis of Dscam diversity in regulating axon guidance in Drosophila mushroom bodies. Neuron 43, 673–686.

